# Magnetized Cellbots to Spatiotemporally Control Differentiation of Human-Induced Pluripotent Stem Cells

**DOI:** 10.1101/2025.03.24.645121

**Authors:** Rashmi P Mohanty, Sudipta Mallick, Peter Crowley, Max Sokolich, Katherine Kiwimagi, Shiva Razavi, Calin Belta, Sambeeta Das, Ron Weiss

## Abstract

Precise spatiotemporal control of gene expression and cellular differentiation is essential for engineering native-like multicellular structures. Current cell differentiation approaches typically rely on externally provided inputs whose effects are not targeted to distinct cells in the appropriate state and hence cannot spatially organize and mature tissue structures as needed. Our work introduces a magnetically controlled microrobot (MR) platform for guiding mammalian cells to desired locations that, combined with synthetic biology, delivers biological signals at precise locations and times, enabling spatiotemporal control of cell-fate decision-making. We use synNotch, a cell-cell contact-based biological signaling that induces relevant gene expression in receivers when the receiver cells contact sender cells through ligand-receptor binding. Magnetically driven MRs are then allowed to be internalized by sender cells, resulting in magnetized sender cellbots. Using a 3-pair orthogonal Helmholtz coil system, we guided magnetized sender cellbots to precise locations in a receiver cell culture, activating desired fluorescent protein expression in target Chinese Hamster Ovary (CHO) receiver cells. Next, we engineered Human-Induced Pluripotent Stem Cells (hiPSC) to function as receivers that can be instructed by senders to differentiate into endothelial cells (ECs) via overexpression of ETV2 (ETS variant transcription factor 2), a master transcriptional regulator of endothelial cell development. Using our magnetic platform, we guided multiple sender cellbots to target locations on a monolayer of hiPSC receivers, resulting in differentiation of receivers into ECs and possible onset of vascular formation. Our approach provides a foundation for the engineering spatial patterns by activating conditional triggers based on MR location and cell state at multiple time points, enabling several applications such as control of organoid architecture.

## 1 INTRODUCTION

High-fidelity tissue engineering, e.g., the ability to grow fully functional organ-like tissue, would enable donor-free organs as well as animal-free drug testing and new disease models, revolutionizing medicine and leading to tremendous societal impact. However, endogenous developmental programs that work remarkably well in natural contexts often fail in non-natural contexts (e.g. outside the context of an embryo). Current approaches that attempt to control cell differentiation in non-native environments often rely on externally-provided inputs (e.g. chemical growth factors) whose effects are not targeted to distinct cells in the appropriate state, and cannot form sophisticated tissue structures. The resulting organoids are size-constrained, limited to a small set of cell types, and do not generally develop fully mature, adult-like tissue. Progress to overcome these obstacles is blocked by the inability to control the spatiotemporal evolution of relevant gene expression patterns. Classical tissue engineering approaches for generating defined structures utilize externally managed construction of cell assemblies such as three-dimensional (3D) bioprinting and hydrogels with considerable control over multicellular organization and initial organoid patterning [1, 2, 3]. However, these top-down approaches are limited to setting the starting conditions for patterning, lacking the dynamic control over the evolution of these structures required to develop native-like tissues.

Alternatively, synthetic biology tools have been developed as bottom-up approaches where synthetic patterning circuits create genetic asymmetries in initially homogenous populations capable of driving multicellular self-organization for creating 2D and 3D patterns toward native-like organoid design. One of the most studied synthetic biology approaches has been the differential expression of cadherin family cell adhesion molecules (CAMs) in mammalian cells for cell sorting [4, 5, 6], 2D and 3D synthetic patterning [7, 8], and morphogenesis [8, 9]. Understanding the molecular mechanism of cell-cell adhesion strength and behavior has been utilized to govern the self-assembly of biological 3D structures that can persist over several days [10, 11]. Over time, arrays of orthogonal synthetic cell adhesion molecules have been engineered to overcome the cross-reactivity of endogenous CAMs and precisely control cell-cell interactions, enabling rationally programmed assembly of multicellular patterns [12, 13, 14]. Similarly, diffusible signaling molecules, called morphogens, are engineered to understand and control patterning *in vitro* [15, 16], *in vivo* [17], and for organoid design [18]. A reaction-diffusion system has also been engineered using a short-range activator and a long-range inhibitor for mammalian cell patterning [19]. Another contact-dependent cell communication pathway, the Delta-Notch pathway, has been used to control cell-type bifurcation [20] and signal propagation [21], which are essential for generating elaborate cell patterns. Further, a synthetic, modular, orthogonal, and combinatorial Notch pathway, the synthetic Notch (synNotch) pathway, has been employed to yield spatial control of diverse cellular behaviors, creating multi-layered self-organizing 2D patterns [22, 23]. For example, a simple synNotch-induced differential cell adhesion molecule expression from a single genotype has resulted in cell fate bifurcation and multi-layer, spatial patterning in spheroids [24]. Recently, synNotch has been integrated into pluripotent stem cells to program differentiation decisions while forming synthetic spatial patterns [25].

Incorporating externally printed cassettes with synthetic biology tools provides additional control over multicellular patterning. Genetically engineered cells capable of establishing intercellular interaction via synthetic biology approaches when seeded on printed cassettes form two-dimensional multicellular patterning with precise spatial control. For example, to form the spatial patterns using CAMs, a nickel-coated surface was functionalized with two synthetic orthogonal CAM-specific proteins for cell adhesion [12]. The specified precise spatial pattern was formed when two distinct cell populations expressing the corresponding orthogonal CAMs were seeded on the functionalized surface. Similarly, surfaces with microcontact-printed functional and orthogonal synNotch ligands have been used to co-transdifferentiate fibroblasts into skeletal muscle and endothelial cell precursors, generating tissues with microscale spatial precision [26]. In another study, synNotch sender and receiver cells were plated in separate chambers of a removable multi-chamber cell culture insert, which was removed once the cells reached confluency, allowing the senders and receivers to make contact, generating a stripe pattern [25]. This method induced contact-mediated neuronal differentiation of receiver pluripotent cells following the synthetic striped pattern. Recently, a multimaterial 3D bioprinting approach resulted in spatially patterned neural tissues when three genetically engineered hiPSC populations were bioprinted to form tri-layer filaments that orthogonally differentiate into neural stem cells, vascular endothelium, and neurons over a global differentiation media condition [27]. Although these approaches create precise spatially controlled programmed patterns, they lack the dynamic control of genetic cues induction, which is essential to guide cell state-based differentiation for forming programmed patterned organoids. Another synthetic biology technique, optogenetics, in which signaling proteins with light-sensitive domains are activated under the control of light, allows dynamic control of gene expression in spatially defined regions. Optogenetically controlled spatiotemporal multicellular morphogenesis has been employed to study developmental biology [28, 29, 30] and to engineer tissue and organoids [31, 32, 33]. However, this approach faces several challenges such as difficulty in light penetration through dense tissue, the need for invasive methods, and addressing the heat generated by light-emitting diodes that can cause tissue damage.

In this work, we introduce a novel convergent platform that combines microrobotics with synthetic biology to obtain precise control in delivering genetic cues to guide cell-state-based differentiation in a spatiotemporal manner. Microrobots (MRs) are micron-sized particles that are promising candidates for different applications due to their versatile size and actuation mechanisms [34, 35]. Specifically, magnetic MRs, which are actuated using an external magnetic field, are extensively studied for biomedical applications because of their high penetration ability and precise controllability [36]. The magnetic platform is advantageous relative to traditional methods like optogenetics for a variety of applications since magnetic fields pass freely through dense biological tissue. Here, we introduce a new type of MR entity, called cellbot, where we embed spherical magnetic MRs inside mammalian cells, enabling robust guidance of the cells to desired locations. Importantly, the MRs are biocompatible and do not affect cellbot function.

We incorporated MRs within sender cells, termed sender cellbots, to communicate with genetically engineered receiver cells via synNotch signaling in a spatiotemporally controlled manner. Using a magnetic control platform, we guided sender cellbots to target locations in receiver cell monolayers, where the sender cellbots activated synNotch in the receivers, providing spatiotemporal control over synNotch activation. In initial validation experiments, we induced expression of a fluorescent protein in spatially targeted CHO receiver cells. In a subsequent experimental setup, we induced expression of ETV2 in sender targeted hiPSC receivers, leading to spatiotemporal control over endothelial cell differentiation (**Fig. 1**). In summary, this study introduces a novel platform combining magnetically controlled MRs with synthetic biology to achieve precise spatiotemporal control over gene expression and cellular differentiation, enabling spatially guided differentiation of hiPSCs into endothelial cells.

**Figure 1:**
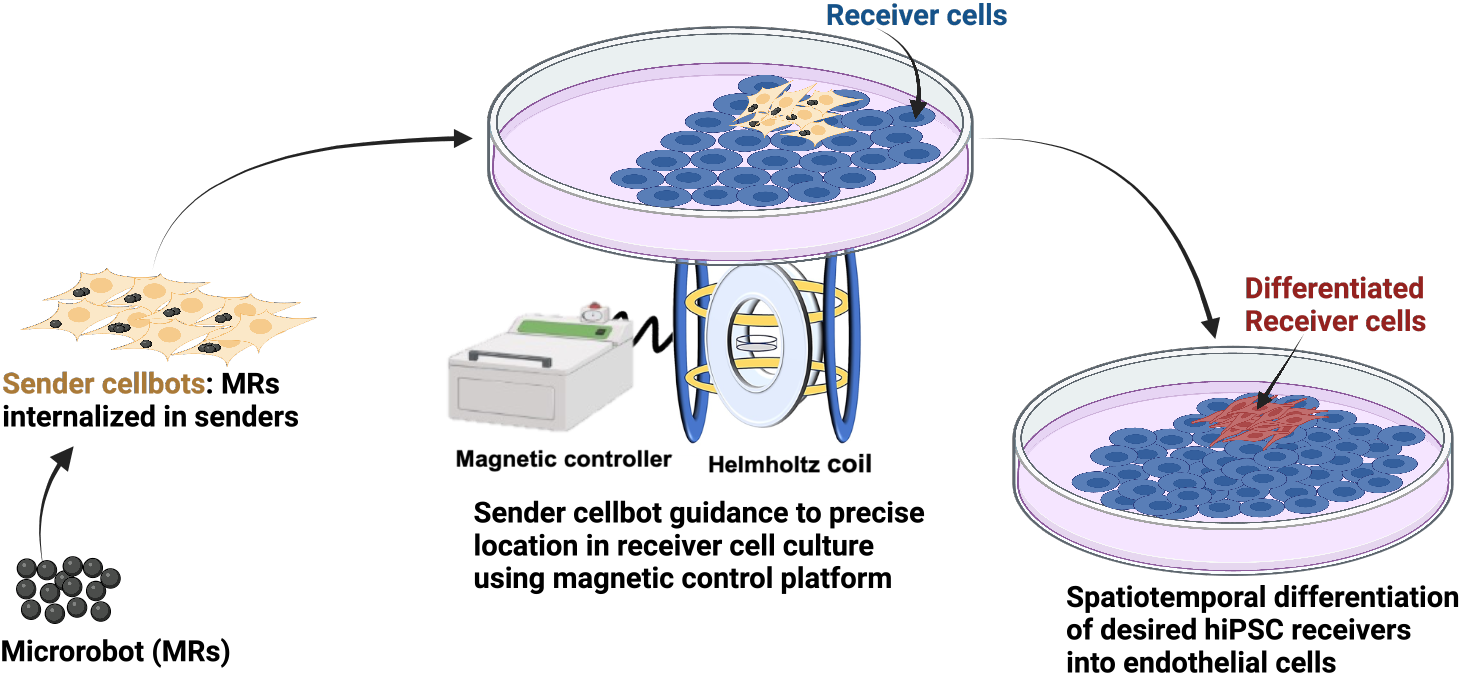
Schematic of MR-guided synNotch-based hiPSC differentiation.

## 2 RESULTS

### 2.1 SynNotch activation in CHO cells

We analyzed synNotch activation dynamics of engineered CHO sender and receiver cells using time-lapse microscopy. The sender cells stably expressed membrane-bound CD19 ligands, and the receiver cells expressed the cognate *α*-CD19 synNotch receptor on their membrane (**Fig. 2a**). When senders come into contact with receiver cells, the ligand-receptor binding leads to proteolytic cleavage of the Notch receptor in the receivers. The intracellular domain fused to the notch receptor contains tTA transcription factor, which is released and translocates into the nucleus due to proteolytic cleavage. tTA binding in the nucleus to a TRE promoter results in expression of a red fluorescent protein, mKate, in activated receiver cells. The sender and receiver cells constitutively expressed a yellow fluorescent protein (YFP) and a blue fluorescent protein (eBFP2), respectively, allowing fluorescence-based cell tracking. We co-cultured sender and receiver cells to quantify synNotch activation, using low sender cell seeding density (10% of total cells) to more efficiently track sender-receiver interactions. Time-lapse images were captured every 2 hours to track the cells and monitor changes in mKate expression in receivers upon synNotch activation. A noticeable overall increase in mKate-positive cells was observed with time (**Fig. 2b & Fig. SI1**) whereas a negative control exhibited no increase in mKate fluorescence (**SI video 1**). The time-lapse images generally indicated that receiver cells expressed mKate after being adjacent to sender cells, and that mKate fluorescent levels in activated receiver cells increased with time (**SI video 2**).

**Figure 2:**
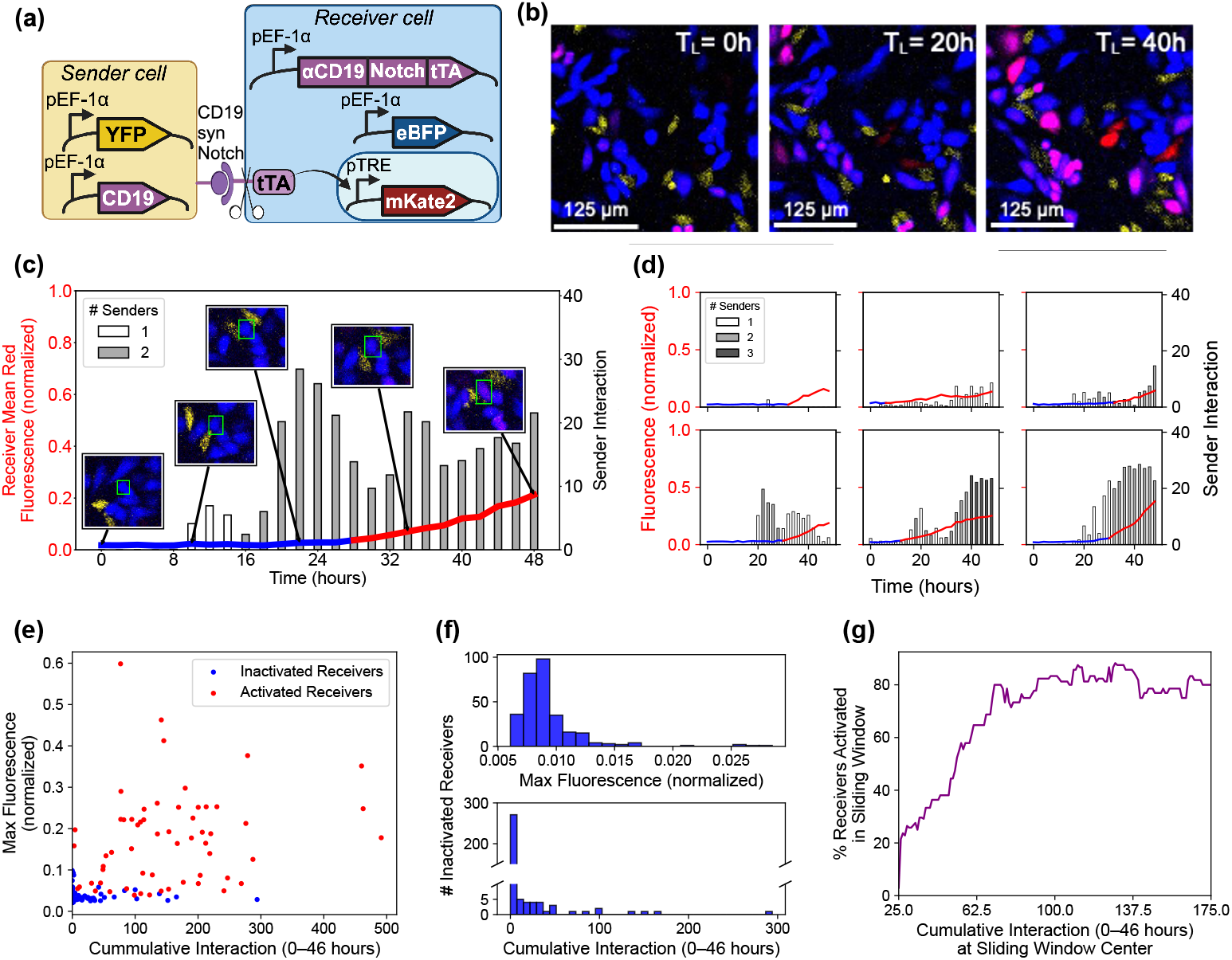
SynNotch activation of CHO receivers by CHO senders. (**a**) Schematics of CD19 synNotch circuit in CHO sender-receiver cells. (**b**) Time-lapse images of synNotch activation in CHO sender and receiver co-culture. Cropped images are shown here, uncropped images are in Figure SI1. Sender cells constitutively express yellow fluorescent protein, receiver cells constitutively express blue fluorescent protein, and synNotch activation in the receiver cells results in expression of a red fluorescent protein. (**c**) A representative activated CHO receiver cell was tracked over a period of 48 hours. The insets show representative fluorescence microscopy images of the tracked receiver cell, surrounded by a green box, at several time points. allowing for visualization of the sender-receiver interactions. (**d**) Each plot in this figure corresponds to a single receiver cell that has been tracked over a period of 48 hours. In Figures (c) and (d), the blue and red line is the cell’s pixelwise mean red fluorescence at each timestep, the indicator of synNotch activation. This line is partitioned into blue and red sections corresponding to pre- and post-observation of activation respectively. Each bar in the graph represents the level of sender interaction with the receiver cell which is calculated based on the length of and distance between sender-receiver interfaces. Bars are grayscale-coded based on number of sender cells in close proximity to the receiver (white=1, light gray=2, and dark gray=3). (**e**) A scatter plot of maximum fluorescence observed at any time in the 48 hours against cumulative interaction from 0 to 46 hours for each receiver cell. (**f**) A set of two histograms of the number of inactivated receivers against maximum fluorescence, and cumulative interaction from 0 to 46 hours. (**g**) A sliding-window plot showing the percent of receivers activated within the sliding window against the cumulative interaction at the center of the sliding window. A window size of 50 and a stride of 1 were used.

Next, we analyzed single-cell synNotch activation dynamics by quantifying individual sender-receiver interactions (as described in Methods) and resulting mKate fluorescence of activated receiver cells over a period of 48 hours. We developed a python-based analysis tool that uses cell tracking data from Cell Profiler [37] to generate graphs that illustrate tracking, fluorescence, and sender interaction information for individual receiver cells in a time series of fluorescent microscopy images (**Fig. 2c**). The graphs also show zoomed-in versions of time series images that focus on a particular receiver cell. We used these images help identify a set of 59 activated receiver cells (out of the 980 total cells identified at 48 hours) that met the following constraints: they were properly tracked by CellProfiler for the entirety of the time series, did not share a common parent cell, did not display mKate fluorescence at time 0, and significantly increased in mKate fluorescence over 48 hours (**Fig. SI2** & **Fig. SI3**). We also identified a set of 300 inactivated receiver cells that were properly tracked by CellProfiler for the entirety of the time series, did not share a common parent cell, did not display mKate fluorescence at time 0, but did not significantly increase in mKate fluorescence over 48 hours (**Fig. SI4**). The activation and interaction data of six representative activated receiver cells chosen from the 59 is displayed in **Fig. 2d**. From analysis of the 59 activated receiver cells, we observed that one sender cell in contact with a receiver cell is sufficient to induce activation. By fitting a ReLU function to each receiver’s fluorescence curve (**Fig. SI2** and **Fig. SI3**), we define the time of activation onset as the ReLU function’s inflection point (transition from 0 to positive slope). We calculated that the average time delay of a receiver cell to activation onset (after interacting with one or more sender cells) is 6.7*±* 4.6 hours after the time of highest sender interaction density (as defined in Methods). Additionlly, we sorted the set of 300 inactivated receivers from low to high interaction and show in **Fig. SI4** a representative set of 20 inactivated receiver cells which span the spectrum of interaction levels. As indicated by the figure, very few of the inactivated receiver cells interacted with senders.

We further analyzed the data in **Fig. 2e, Fig. 2f, Fig. 2g, Fig. SI5**, and **Table SI1** to determine the relationship between sender-receiver interaction and resulting receiver mKate fluorescence. In **Fig. 2e**, we plot each receiver’s cumulative interaction over the first 46 hours against its maximum recorded fluorescence. We do not consider the interaction at the final timestep (48 hours) both because this does not influence the fluorescence over the 48 hours, and because one standard deviation beneath the average delay time in activation *≈* 2 hours, which makes 46 hours the latest time that sender interaction could have activated a receiver. In **Fig. 2f**, we illustrate the high number of inactivated receivers clustered around the origin in **Fig. 2e** by showing the distribution of cumulative interaction and maximum fluorescence for the inactivated receivers. In **Fig. 2g**, we show that the likelihood of activation increases with increasing cumulative interaction. In **Fig. SI5** we show the Receiver-Operating Characteristic (ROC) curve for predicting whether a receiver is activated or inactivated based on a cumulative interaction threshold. **Table SI1** shows the confusion matrix of using a value of 30 cumulative interaction as the prediction threshold. In this case, 93% of the activated receivers were measured to have cumulative interaction greater than or equal to this threshold, and 95% of inactivated receivers were measured to have cumulative interaction lower than this threshold.

We expand our analysis by studying the correlation between sender interaction and the derivative of mKate fluorescence. In **Fig. SI6a**, we generate a scatter plot of each receiver’s slope of mKate fluorescence from the ReLU curve-fitting versus cumulative interaction. Note that all inactivated receivers have a ReLU slope of zero. In **Fig. SI6b**, we illustrate the high number of inactivated receivers clustered around the origin in **Fig. SI6a** by showing the distribution of cumulative interaction for the inactivated receivers. In **Fig. SI6c**, we show a generally positive correlation between cumulative interaction and fluorescence derivative.

### 2.2 Electromagnetic control platform for precise delivery of magnetized sender cellbots

We developed magnetized sender cellbots, which are sender cells containing magnetic microrobots (MRs), to deliver synNotch signaling to precise locations in a receiver cell culture using our magnetic control platform. MRs are nickel-coated silica spheres of 4.8 *µ*m diameter, where nickel coating on the silica microspheres was performed to magnetize the microspheres using physical vapor deposition (PVD) [38]. We set the thickness of the nickel coating to 100 nm using a PVD evaporation machine, as this thickness was found to be sufficient to drive sender cells using our magnetic setup. SEM images confirm that nickel deposition was effectively achieved on the upper half of the microspheres (**Fig. 3a**), as expected with the PVD method. Successful nickel coating appears as dark patches on the microspheres under optical microscopes (**Fig. 3a**). When incubated with sender cells in culture, the MRs were internalized into the sender cells within 24 hours (**Fig. 3b**). These magnetized sender cellbots maintained the distinct spindle-shaped morphology of CHO cells in their typical culture conditions (**Fig. SI7a**) and exhibited less than 10% decrease in cell viability compared to sender cells without MR incubation (**Fig. SI7b**). Upon cell division, sender cellbots pass the MR to one of the daughter cells (**Fig. SI7c**), while initiating the formation of a micro-colony, indicating that MR-loading does not significantly hinder cell proliferation.

**Figure 3:**
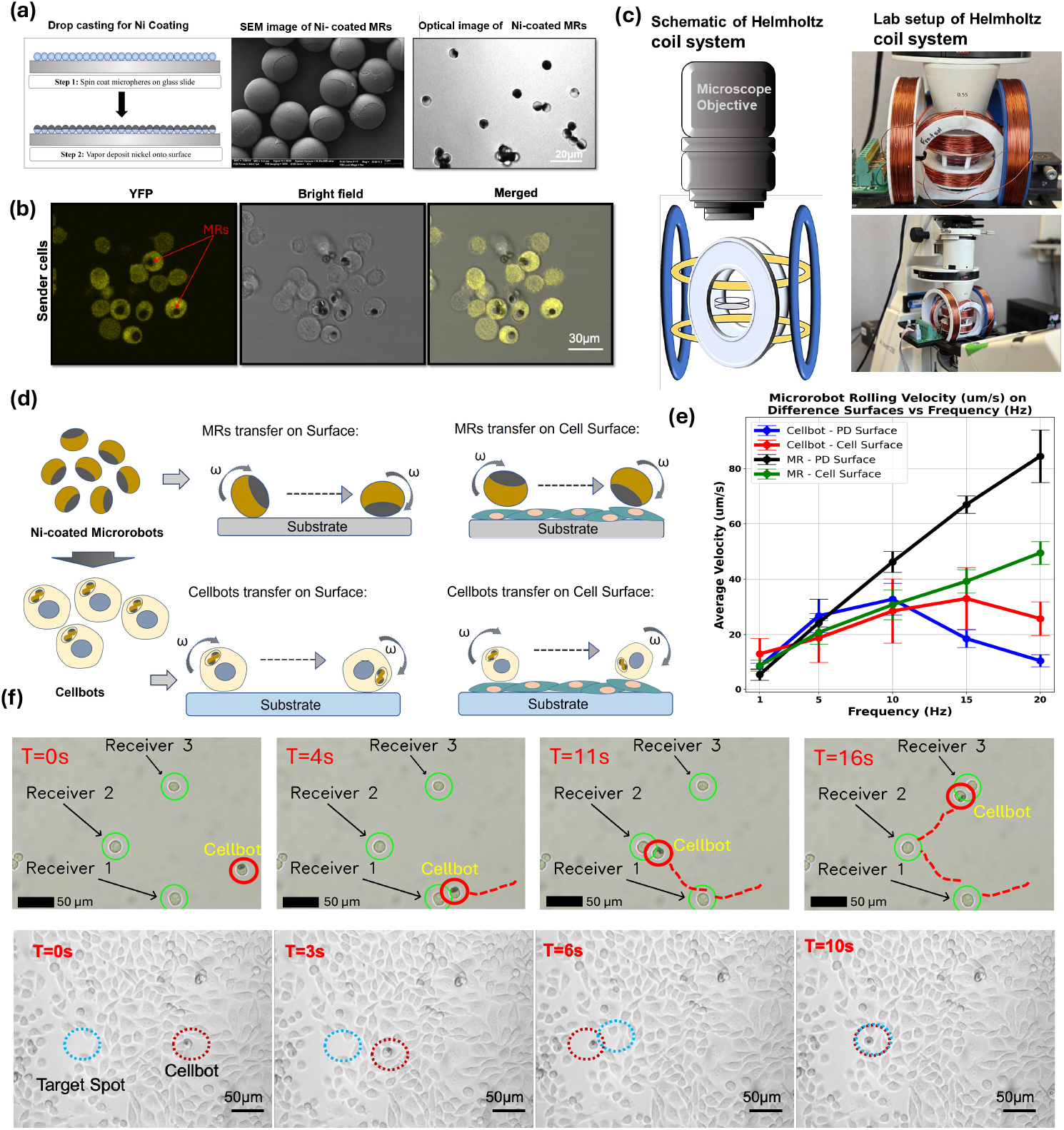
Magnetic control platform for MR and Cellbot control. (**a**) Illustration of Microrobot fabrication followed by Scanning Electron Microscopy (SEM) and optical microscopy images of Ni-coated MRs. (**b**) Confocal images of Sender cellobots showing internalized MRs. (**c**) Schematic representation of Helmholtz Coil system and pictures of experimental lab setup. (**d**) Illustration of rolling MRs and cellbots on Petri dishes with glass surface and on a surface populated by cells. (**e**) Graph depicting speed vs frequency of MR and sender cellbot movement on the two different surfaces. (**f**) Time-lapse images showing targeted delivery of magnetically-driven sender cellbots to desired receiver cells on glass surface (upper panel) and cell surface (lower panel). Scale bar = 50 *µ*m

We used an electromagnetic manipulation platform to navigate the MRs and sender cellbots to user-defined locations. The electromagnetic manipulation platform consists of a 3D Helmholtz coil system, a microcontroller, power electronics, and custom control software necessary to generate rotating magnetic fields in the workspace (**Fig. 3c**) [39]. The fields generated by the system were measured with a tesla meter to be approximately 3 mT. On a given surface, these rotating magnetic fields guide the rolling motion of MRs, and hence magnetized sender cellbots, in any desired 2D direction. To generate rotating magnetic fields using a Helmholtz coil system, sinusoidal signals are sent to each pair of coils as described in the SI. Accurate adjustment of the rotation axis of the field enables fine-grain control over MR and sender cellbot locomotion.

Using our magnetic control platform, we assessed the relationship between magnetic frequency and speed of guided cellular motion on both glass surface as well as cell monolayer (schematic representation in **Fig. 3d**). We first performed a control experiment with just controlling the rolling motion of MRs on a glass petri dish and on a cell surface at 5, 10, 15, and 20 Hz rotating magnetic field. The result indicate that the MR movement can be guided effectively on both surfaces and MR speed increases with rotating field frequency (**Videos 3 and 4**). We observed slower MR movement over the cell surface in comparison to glass surface. Similar rolling experiments were conducted using sender cellbots on both glass and cell surfaces (**Videos 5 and 6**). We analyzed the videos to determine the measured speed of MRs and sender cellbots as a function of rotating magnetic field frequency on both surfaces **Fig. 3e**. A linear increase in speed was observed for MRs on both glass and cell surfaces whereas the speed for cellbots increases up to a certain frequency and then starts to decrease. This frequency is called the step-out frequency of rotating objects above which the applied magnetic torque is not strong enough to keep the rotating object synchronized with the field [40] (described in SI).

Then, we evaluated the effect of magnetic actuation on cell viability to ensure that it does not elicit significant toxicity. We exposed the cellbots and cells without MRs to a 10 Hz magnetic field for 15 mins and then incubated the cells for 24 hrs. As shown in Fig. SI8a, both cellbots and cells without MRs were multiplying and their morphology was intact after magnetic actuation when compared to control cells without magnetic actuation. Flow cytometry data revealed that while the MR loading results in a slight decrease in cell viability as discussed above (**Fig. SI7b**), the impact of actuation appears to be almost negligible with the measured 2% further decrease in cell viability (**Fig. SI8b**). **Figure 3f** shows the ability of the electromagnetic manipulation system to guide sender cellbots to receiver cells. Using the gaming controller joystick described in SI, we guided movement of a sender cellbot to each of three receiver cells observed in the same field of view (**Figure 3f, Video 7**). It took approximately 16 seconds to move the sender cellbot in a path that brought it in close proximity to these three cells, traveling a rough distance of 300 µm. The bottom panel in **Figure 3f** shows the successful delivery of a sender cellbot to a target position, over a surface densely populated by receiver cells. In this case, it took the sender cellbot around 10 seconds to travel approximately 150 µm.

### 2.3 MR-guided synNotch activation in CHO cells

Next, we evaluated whether loading the MRs into the senders to create sender cellbots interferes with the ability to communicate with receiver cells. We co-cultured senders and sender cellbots with receiver cells and observed that both senders were able to activate synNotch signaling in receiver cells (**Fig. 4a**). As expected, WT cells (cells w/o CD19 ligand) did not activate synNotch signaling. We next detached the cells from the co-culture plates using Versene and quantified the mKate fluorescence intensity of receivers from both cultures. We found that receiver mKate fluorescence is the same when co-cultured with either sender cellbots or sender cells without MRs (**Fig. 4b**). We then magnetically guided sender cellbots to precise locations in a receiver cell culture (**Fig. 4c, Fig. SI9, Video 9**). **Fig. 4d** shows an example of time series images illustrating the increase in mKate expression of two receivers over time. We performed the same single-cell synNotch analysis as described in **Fig. 2c** & **Fig. 2d** to quantify one of the two sender-receiver interactions and receiver activations (**Fig. 4e**), and noted that receiver activation was observed after 12h of interactions with senders.

**Figure 4:**
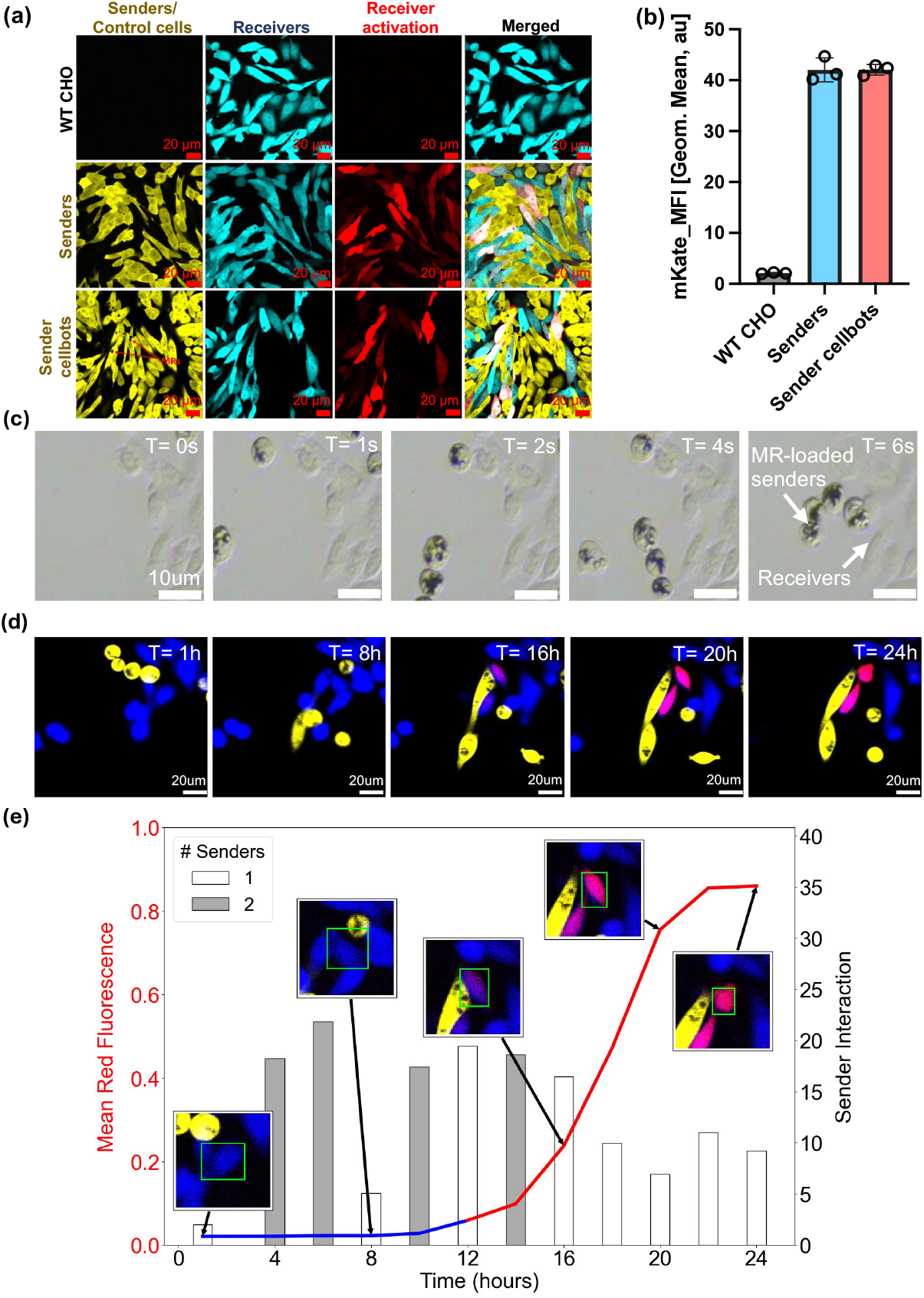
MR-guided synNotch activation in CHO cells. (**a**) Time-lapse images showing synNotch activation when MR-loaded CHO senders are cultured with CHO receivers. (**b**) Quantification showing MR-loading does not impact mKate expression (or synNotch activation). (**c**) Sender cellbots were delivered to target receiver cells in a CHO receiver culture. (**d**) Time-lapse images showing synNotch activation when the CHO senders cellbots maintained contact with target CHO receivers. Cropped images of c and d are shown here, uncropped images are in Fig. SI9. (**e**) A single receiver CHO cell (blue) has been tracked over a period of 24 hours. The insets show the fluorescence microscopy images from Figure (d) zoomed in on the tracked receiver (boxed in green), allowing for visualization of the sender-receiver interactions.

### 2.4 SynNotch-based hiPSC differentiation into endothelial cells

Next, we engineered hiPSC receiver cells that can differentiate into endothelial cells (ECs) by overexpressing ETV2 upon synNotch induction. ETV2 (ETS variant transcription factor 2) is a master transcriptional regulator of endothelial cell development that has been used to guide the differentiation of hiPSCs into ECs [41, 42]. We genetically modified the synNotch receivers to express ETV2 upon activation (**Fig. 5a**). Once receivers are in contact with senders, the ligand-receptor binding event leads to proteolytic cleavage of the Notch receptor, releasing tTA, which then translocates into the nucleus to activate the TRE3G promoter, expressing ETV2 and mKO2 in the receiver cell (**Fig. 5a**).

**Figure 5:**
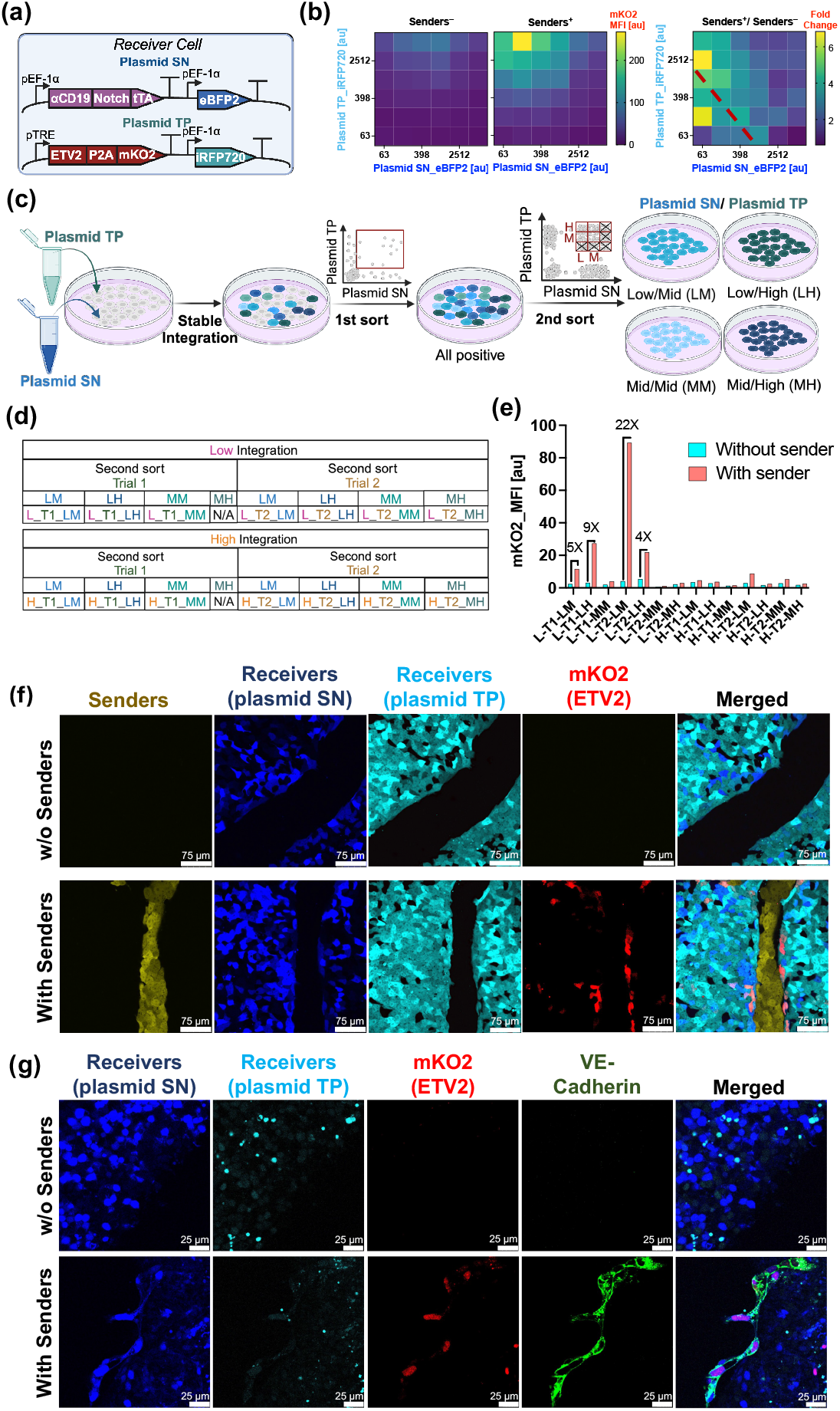
ETV2-expressing hiPSC cells differentiate into endothelial cells when co-cultured with HEK sender cells. (**a**) Genetic circuit for synNotch based ETV2 expression. (**b**) Transient polytransfection of genetic circuit indicates importance of stoichiometry between the two receiver cell plasmids. (**c**) Schematic of stable integration of plasmids in hiPSC receivers and cell sorting strategy to select the best performing receiver cell subpopulation. (**d**) Measured receiver cell activation for 14 polyclonal cell lines and (**e**) selected the best performing hiPSC receiver (L T2 LM) for further study. (**f**) Confocal imaging shows synNotch activation-based ETV2 expression in hiPSC receivers when co-cultured with HEK senders (**g**) Immunofluorescence verifies presence of endothelial cell (EC) specific marker, VE-Cadherin, that indicates hiPSC receiver cell differentiation into ECs upon synNotch activation.

The receiver circuit was divided into two plasmids to facilitate DNA construction and testing of optimal stoichiometry of circuit components (**Fig. 5a**). We introduced the two receiver cell plasmids into HEK293FT cells using poly-transfection [43], a one-pot transfection strategy that enables uncorrelated transfection of plasmids in cells, generating cell populations with different plasmid stoichiometries. HEK stable senders (with CD19 ligand) were generated and were co-cultured with the transfected receiver cells. A co-culture of the transfected receiver cells with WT HEK cells (without CD19 ligand) was used as a negative control. Constituitive expression of eBFP2 and iRFP720 fluorescence markers from the plasmids were used to estimate plasmid stoichiometries in the individual cells. SynNotch-based ETV2 expression was tracked with mKO2 reporter, providing for the generation of multi input heat maps for receiver activation (**Fig. 5b**). In co-culture with senders, maximal receiver cell activation was observed for low-to-medium levels of synNotch receptor expression (Plasmid SN) and high levels of target promoter (Plasmid TP). High fold receiver activation relative to negative control was observed for low-to-medium Notch receptor expression (Plasmid SN) coupled with medium/low and medium/high levels of target promoter (Plasmid TP) (**Fig. 5b**).

Next, we generated hiPSC receiver cell lines by stably integrating the two plasmids into WT hiPSCs. We aimed to generate both low and high integration copy numbers by varying the transfected plasmid DNA amounts. After integration, we sorted cells positive for both eBFP2 and iRFP720 in the first round of sorting (**Fig. 5c**). Once we recovered enough double-positive cells from the first sort, we used the cells for a second round of sorting. Here, we divided the double-positive cells into nine subpopulations based on low (L), medium (M), or high (H) levels of eBFP2 and iRFP720 expression. Based on the transient transfection heatmap matrix results (**Fig. 5b**), we focused on collecting four target subpopulations from each of low and high transfection amounts: low/medium Plasmid SN and medium/high Plasmid TR (**Fig. 5c**). We performed the second round of sorting twice for both integrations, collecting a total of 16 cell lines. However, we were not able to recover two of the cell lines, yielding a 14-cell line library (**Fig. 5d**). We screened the 14 cell lines for their synNotch activation potential by co-culturing them with monoclonal HEK sender cells (**Fig. 5e**). The top four performers all came from the low integration batch and had an observed low copy number for Plasmid SN (**Fig. 5e**). The top performer, L T2 LM, exhibited 22-fold synNotch activation when co-cultured with senders as compared to without sender cells. This cell line was selected for the remainder of the studies, and is henceforth referred to as hiPSC receiver cells.

In co-culture, HEK senders and hiPSC receivers self-organized to form distinct colonies. After two days of co-culture, we observed receiver cells, mostly at the interface of the sender-receiver colonies, expressing mKO2 fluorescent protein (**Fig. 5f**), and suggesting synNotch-based co-expression of ETV2. In contrast, in a co-culture with WT HEK cells (cells without the CD19 ligand) as a negative control, we did not observe mKO2 (or ETV2) expression (**Fig. 5f**). We continued the sender-receiver co-culture in mTeSR pluripotency maintaining media to demonstrate the differentiation of ETV2 over-expressing hiPSC receivers into endothelial cells. After 8 days of co-culture, we quantified transcripts of ETV2, EC-specific markers, and pluripotency markers using RT-qPCR (**Fig. SI10**). Overall, we observed significantly higher expression of ETV2 (**Fig. SI10a**) and EC-specific markers, such as vascular endothelial cadherin (VE-Cadherin) (**Fig. SI10b**), platelet endothelial cell adhesion molecule 1 (PECAM-1 or CD31) (**Fig. SI10c**), and CD34 (**Fig. SI10d**), in sender-receiver co-culture as compared to receivers co-cultured with WT cells. We also observed a significant reduction in pluripotency markers octamer-binding transcription factor 4 (OCT4) (**Fig. SI10e**) and Nanog (**Fig. SI10f**), indicating differentiation of hiPSC receivers into endothelial cells triggered by synNotch activation. Using immunofluorescence, we also confirmed VE-Cadherin expression in ETV2-expressing hiPSC receivers, but not in a co-culture without sender cells (**Fig. 5g**).

### 2.5 MR-guided hiPSC differentiation into endothelial cells

Finally, we aimed to deliver a large number of sender cellbots to a broad target region in an hiPSC receiver colony, triggering nearby hiPSC receivers to differentiate into endothelial cells (ECs). We coated half of a 35-mm dish with matrigel, allowing hiPSCs to grow only on the matrigel-coated side (**Fig. 6a**). Then, we added approximately ten thousand sender cellbots on the non-matrigel coated side of the dish and guided multiple magnetized HEK sender cellbots to target locations of hiPSC culture using our magnetic-control platform (**Fig. 6b**). We did not observe appreciable mKO2 expression in hiPSC receivers just after sender cellbot guided movement to the target region (**Fig. 6c**). We then maintained the co-culture in an incubation chamber and acquired fluorescence images at several time points. We observed mKO2 (and implied ETV2) expression in hiPSC receivers located in the target region within 24 hours (**Fig. SI11a**) and continued monitoring mKO2 expression for several additional days (**Fig. 6d & SI11b**). After eight days of incubation, we immunostained the co-culture and imaged a large band of the matrigel coated side of the plate for all relevant fluorescent channels. The fluorescence imaging suggests that the sender cellbots resided mostly in the targeted region even after eight days of co-culture, resulting in hiPSC differentiation and possible onset of vascular formation mostly in the target region (**Fig. 6e, Fig. SI12, & Fig. SI13**), but not appreciably in the untargeted locations that we examined (**Fig. SI14**).

**Figure 6:**
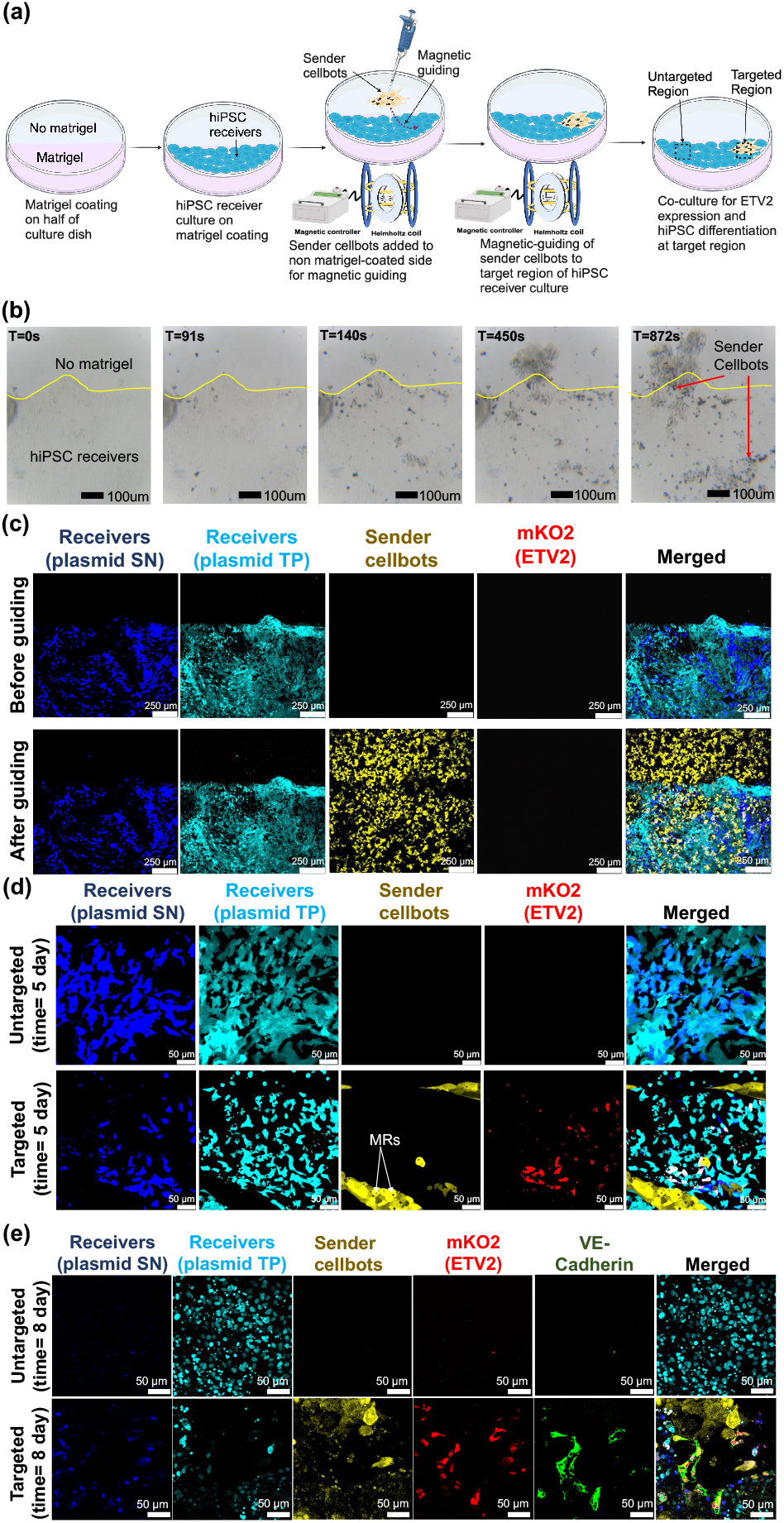
MR-guided hiPSC differentiation at target location. (**a**) Schematics of sender cellbot guidance to target region using magnetic control platform. (**b**) Time-lapse bright-field images of sender cellbot guidance to a target region of hiPSC receivers. (**c**) Fluorescence images of the target region before and after sender cellbot guiding. (**d**) SynNotch activation-based mKO2 (and ETV2) expression in hiPSC receivers in the target region after 16 and 24 hours. Cropped images are shown here, uncropped images are in Fig. SI10. (**e**) Fluorescence imaging of a large band shows significant sender cells resided in the targeted region after eight days, resulting in VE-cadherin expression in hiPSC receivers in the targeted region.

## 3 Discussion

Precise spatiotemporal control of gene expression is essential for developing native-like or customized multicellular structures. Our work introduces a magnetically guided micro-robotic platform that can deliver biological signals at precise locations to activate genetic cues for cell-state perturbations. We use synNotch, a cell-cell contact-based biological signaling mechanism that induces desired gene expression in receiver cells upon ligand-receptor binding with sender cells. To enable precise guidance of sender cell localization, we internalized magnetic MRs in sender cells, resulting in magnetized sender cellbots. Using a magnetic platform, we guided the sender cellbots to precise locations in a receiver cell culture to activate desired gene expression in target CHO and hiPSC receiver cells. In CHO cells, we expressed a desired fluorescent protein in target receivers. In hiPSCs, we overexpressed ETV2 in receivers located in the target region, leading them to differentiate into endothelial cells, while hiPSCs in the untargeted region remained undifferentiated. Magnetic-based actuation allowed sender cellbot guidance and biological signal delivery to user-defined locations, resulting in hiPSC differentiation in a spatiotemporal controlled manner.

Following previously published works [22, 44], we engineered CD19 and *α*CD19/tTA synNotch expressing CHO sender and receiver cells, respectively (**Fig. 2**). In bulk experiments of 1:1 mouse embryonic stem cell sender-receiver co-culture, others have observed synNotch activation kinetics with a 5–6 hours lag time of reporter protein expression after the initial sender-receiver contact occurred [25]. Studies also reported that activation pulses as short as 2 hours are sufficient to induce significant levels of reporter protein expression, and the protein expression can continue for several hours even after the sender-receiver contact is lost [22, 25]. In our single cell synNotch activation dynamics assays, we observed that contact with one sender cell is sufficient to activate synNotch in receiver cells (**Fig. SI2 & Fig. SI3**). Once an interaction happens, we approximated the average delay for observing synNotch activation to be 6.7 *±* 4.6 hours. Fluorescence protein expression often continued to increase during the 48 hour experiment even after apparent discontinuance of cell-cell contact.

To enable guided control over sender cellbot localization, we loaded the Nickel-coated silica MRs into the cells by co-incubation (**Fig. 3**). MR-loading did not impact synNotch activation functionality of sender cellbots or the corresponding receiver cell activation (**Fig. 4a & 4b**). Upon guiding sender cellbots to precise receiver cells using the magnetic control platform (**Fig. 4c**), we observed synNotch activation in target receiver cells within 12 hours of co-culture (**Fig. 4d**). Time-lapse microscopy validated that sender cellbots contained MRs during the entire study period of 26 hours. The measured speed of the sender cellbots of 25 um/sec at a 3Hz rotating magnetic field frequency is sufficiently fast such that it should not induce synNotch activated gene expression in receiver cells along it’s route to the target cells. Also, for all synNotch activated receivers we always found corresponding sender cells that were likely to be the ones that initiated the activation.

Spatiotemporal control over vascular formation has been a significant challenge for engineering native-like organoids. Toward the formation of vascularization in a spatiotemporally controlled manner, we used our MR-guiding platform to activate synNotch signaling at precise locations in an hiPSC culture and differentiate the targeted hiPSCs into endothelial cells. Previous studies reported the integration of synNotch circuits in Rosa26 landing pad sites of mouse ESCs to generate clonal receiver cell populations to overexpress Neurog1 transcription factors for synNotch-based neuronal differentiation [25]. In comparison, we integrated synNotch receiver gene circuits in hiPSCs using a poly-piggyBac transposon system to overexpress the transcription factor ETV2 upon synNotch activation to lead the receivers toward endothelial cell fate. Poly-PiggyBac integration allowed optimization of stoichiometry of the sensing and responding elements of the receiver cell circuits (**Fig. 5c**). The sensing element contains transcription factor tTA that, upon synNotch-based proteolytic cleavage, bound to TRE promoter to express ETV2 (**Fig. 5a**). We found that a low tTA to TRE promoter ratio is optimal for obtaining a high fold-change in synNotch activation (**Fig. 5b**). This observation is consistent with our previously published work which reported that lower levels of synNotch receptors improved the performance of receiver cells primarily because of the decrease in leaky output from the uninduced receiver cell populations [44]. The combination of transient transfection analysis (**Fig. 5b**) and stable piggybac integration using one-pot poly-transfection [43] allowed us to efficiently sort for cells with optimal stably integrated genetic circuit stoichiometry (**Fig. 5c, 5d & 5e**). Using this methodology we obtained a polyclonal receiver cell population exhibiting significant fold change (22-fold) in synNotch activation.

In this paper, we discussed both CHO and HEK senders. CHO cells were used primarily for the initial sets of experiments because they tend to remain as single cells more often than HEK cells, and as such their single cell communication behavior is easier to quantify. We opted to use HEK sender cells for hiPSC experiments due to their increased aggregation phenotype, which helps increased efficiency of guided hiPSC differentiation. In co-culture, HEK and hiPSC cells self-assembled to form distinct colonies due to the types of cell adhesion molecules expressed on their surface [45, 46]. Hence, in a HEK/hiPSC sender-receiver co-culture, we observed synNotch activation mainly at the interface of different cell clusters (**Fig. 5f**). Once we confirmed cell-cell communication with fluorescent reporters, we focused on analysis of hiPSC overexpression of ETV2 upon synNotch activation (**Fig. SI10a**). Previous studies have demonstrated that overexpression of transcription factors in programmed hiPSCs overrides pluripotency maintaining media or media-driven differentiation [27]. Hence, we maintained the synNotch-based ETV2 overexpressing hiPSCs that were co-cultured with HEK sender cells in pluripotent stem cell maintenance media for eight days while monitoring hiPSC guided differentiation into endothelial cells. We verified presence of EC-specific markers such as VE-cadherin (**Fig. 5g & Fig. SI10b**), CD31, and CD34 (**Fig. SI10c & d**), suggesting ETV2 overexpression guided receiver cells toward endothelial cell state. We did not observe late endothelial cell markers, such as Von Willebrand factor (vWF), within the eight days of differentiation. Other studies have observed vWF after similar duration of Dox-induced ETV2-driven differentiation of hiPSCs [27, 47]. In comparison to these studies, we wanted to ensure that only the desired hiPSCs within the population differentiated into ECs. Hence, we cultured all the cells in pluripotency maintaining media, while other studies supplemented growth factors in their cultures, likely leading to a faster response.

Using our MR-control platform, we guided multiple HEK sender cellbots to desired locations of hiPSC receiver culture for the induction of spatiotemporal ETV2-based differentiation of targeted receivers into endothelial cells (**Fig. 6**). The constantly proliferating sender and receiver cells over the study period of eight days confirmed that the MRs and magnetic field actuation minimally imparted cytotoxic effects on HEK and hiPSCs. For future applications with hiPSC differentiation, we will engineer sender cells to contain additional safety switches so that they can be eliminated once the desired biological signaling is activated in receivers. With our current micro-robotic setup, we could alter the cell fate of receivers in the targeted region while the hiPSCs in the untargeted region remain unchanged (**Fig. 6e, Fig. SI12, Fig. SI13 & Fig. SI14**).

In conclusion, we demonstrated both coarse-grained and fine-grained control over sender cellbot localization and resulting hiPSC to endothelial cell differentiation. Achieving more precise multilocation spatiotemporal patterning with magnetic microrobots will require additional advances. In the future, we will explore alternative actuation approaches such as combining rotating magnetic fields with magnetostatic selection fields, developing closed-loop control strategies, and investigating alternative actuation methods like sound-powered microrobots. Striving towards improved control to deliver biological signaling at a single-cell level precision in receiver cell cultures, our work, a combination of microrobotics and synthetic biology, provides a unique and novel platform to alter cell fate in a spatiotemporal manner. The MR-guiding platform enables spatiotemporal control and dynamic perturbations over gene expression and MR-loaded sender cellbots could be genetically engineered to incorporate user-defined functionalities. Such sender cellbots would support continuous, bidirectional communication with receiver cells, thereby enabling sophisticated spatiotemporal control of gene expression. We anticipate that our findings will pave way for next generation organ-like tissue engineering and organoid formation for regenerative medicine, disease modeling and donor-free organ transplant applications.

## 4 Methods

### 4.1 Cell culture

Adherent Chinese Hamster Ovarian (CHO-K1, wild type) cells were originally obtained from the American Type Culture Collection (ATCC). CHO-K1 cells with a monoallelic integrated landing pad (LP) in the putative Rosa26 locus [48] (CHO-K1-LP) were genetically engineered to generate CHO synNotch sender and receiver cell lines. All CHO cells were maintained in Dulbecco’s Modified Eagle Medium/Nutrient Mixture F-12 (DMEM/F-12, Gibco) supplemented with 10% fetal bovine serum (FBS) (Cellgro). HEK293FT cells (wild type) were purchased from Invitrogen. HEK293FT-LP cells [48] were used to generate HEK synNotch sender cell lines. All HEK cells were cultured in DMEM (Corning) supplemented with 10% FBS. Alstem 11 hiPS cells were purchased from ALSTEM and were cultured in mTeSR Plus media (StemCell Technologies). For hiPSC cultures, we coated tissue culture plates with Matrigel (Corning), diluted according to the manufacturer’s instructions in ice-cold DMEM/F-12. Matrigel-coated plates were maintained at 37°C for 30 min before seeding the hiPSCs. Cells were incubated with Gentle Cell Dissociation Reagent (GCD, StemCell Technologies) for 6 min at room temperature to be passaged as aggregates. Then, the GCD was aspirated, and cells were mechanically dissociated with cell scraper (Corning) in fresh mTeSR. Dissociated cells were gently resuspended in fresh mTeSR so as not to dissociate the hiPSCs into single cells and were replated. We passaged hiPSCs as single cells by incubating them in GCD for 10 min at 37°C, followed by re-suspending the detached cells in fresh mTeSR and centrifuging the suspension at 1000 RPM for 3 minutes. The cell pellet was then resuspended in fresh mTeSR medium supplemented with 10*µ*M Y-27632 (ROCK inhibitor, Tocris Biosciences) before seeding. The culture was replaced with fresh mTeSR plus within 16 hours. All cell lines used in the study were grown in a humidified incubator at 37°C with 5% CO2 and passaged every 2–3 days. All experiments were performed after third cell passage. All the cell lines were tested for mycoplasma.

### 4.2 Plasmid construction

The basic sender and receiver synNotch genetic components were obtained from a previously published work [44]. A complete list of plasmid constructs used in the study can be found in **Table SI2**.

#### Lenti constructs

The plasmid encoding membrane-bound CD19 ligand (pKK499) was constructed in a lentiviral transfer backbone to engineer CHO and HEK sender cells. Plasmids, pKK493 and pKK562, were constructed in lentiviral transfer vectors to engineer CHO receiver cell. pKK493 plasmid encodes synCD19 ScFV fragment, Notch core, and tTA transcription factor to be expressed on the transmembrane. pKK562 plasmid contains, TRE promoter expressing mKate. Another plasmid, pKK589, was constructed in a landing pad payload vector to encode the reporter EBFP2 protein and the puromycin antibiotic resistance gene.

#### MoClo construct

To engineer hiPSC SynNotch receiver cells, we constructed plasmids based on a modified hierarchical MoClo system [49]. Genetic elements were first cloned into corresponding level 0 (L0) destination vectors; insulators (pL0.I), promoters (pL0.P), 5^*′*^ UTR (pL0.5^*′*^), gene (pL0.G), 3^*′*^ UTR (pL0.3^*′*^) and terminator (pL0.T). Then, the genetic elements from L0s were assembled into L1 positioning expression vectors for each transcriptional unit (TU), with ST1-2 for position 1, ST2-3 for position 2, and ST3-4X for position 3. Finally, the L1 vectors were assembled by Golden Gate cloning using SapI into an L2 PiggyBac transposon vector, which was used either for transfection-based assays or for stably integrating genetic elements into cell chromosome. We constructed two L2 plasmids (pRM105 and pRM106) based on MoClo assembly. pRM106 (or Plasmid SN) encodes synNotch sensing element in the first TU and EBFP2 fluorescent protein reporter in the second TU. pRM105 (or Plasmid TP) encodes synNotch responding elements, TRE3G promoter activating co-expression of ETV2 and mKO2 reporter in the first TU. The second TU encodes constitutive expression of iRFP720 fluorescent reporter. ETV2 in the pL0.G backbone was obtained from our previously published work [50].

### 4.3 Genomic Integration

#### CHO & HEK senders-receiver cell lines Lentiviral Production

Lentivirus was created using payloads built into pLV R2R2 GTW3 backbone. These payloads were transiently cotransfected with pCMV-dR8.2 dvpr (Addgene, #8455) and pCMV-VSV-G (Addgene, #8454) in HEK293FT cells. Transfected media was replaced with fresh media after 24 hours of transfection. After 48 hours, virus was harvested by filtering virus-contained media through a 0.45 *µ*m polyethersulfone (PES) membrane. Virus was used the same day of collection.

#### Lentiviral transduction

A 24-well plate of HEK293FT-LP or CHO-K1-LP cells at 50% confluency were incubated with filtered virus at 3 different concentrations, 1:4, 1:12 & 1:36 virus:media, in a total volume of 400 *µ*l DMEM media per well supplemented with 10% FBS, 1% non-essential amino acid, and 10 mg/ml Polybrene. Virus-containing media was replaced with fresh media after 24 hours. The resulting polyclonal populations were expanded and underwent two passages before further analysis.

#### Landing Pad Integration

Landing pad cell lines contain the constitutive expression of YFP that gets knocked out during the integration of payload vectors, as previously published [48]. To utilize this technology, cells were transiently co-transfected in a 24-well plate with the payload vector and vector containing a construct expressing BxB1. When cells reached 90% confluency, they were transferred to a 6-well plate. When cells reached 50% confluency in the 6-well plate, 10,000x (10mg/ml) Puromycin stock was used to create 2x puromycin-containing media. Cells were grown in Puromycin-containing media with daily media changes until reached 90% confluency. Then the cells were transferred to a 100 mm dish, and puromycin was removed from future media changes.

#### Sorting

For HEK and CHO sender cell lines, cells were sorted after five days of lentiviral transduction and after two passages. Cells were live stained with HA Tag antibody (Invitrogen, #26183) and were then single cell sorted for florescence into 96-well plates. Monoclonal cells were then screened for functionality through co-culture with transiently transfected receiver cells. For CHO receiver cell, monoclonal sort was done after lenti transduction. Five days after transduction and after two passages, cells were co-cultured with transiently transfected CHO sender cells or WT CHO cells for 2 days. Co-cultures with WT CHO cells were used to set the mKate expression gate by placing the sorting gate just higher than the highest leaky expression cell. Co-cultures with sender cells were used to sort single cells into a 96-well plate for cells with expression higher than the mKate expression measured with the WT co-culture. Monoclonal populations were then screened for functionality through co-culture with transiently transfected sender cells. One of the high-functioning clones was chosen for landing pad integration to add EBFP2 constitutive expression.

#### hiPSC receiver cell line

##### PiggyBac Integration

PiggyBac integration [51], in combination with poly-transfection [43] was used for the genomic integration of MoClo plasmid constructs to engineer hiPSC receiver cell lines, which we call poly-PiggyBac integration strategy. We acquired both a low and a high DNA copy number after chromosomal integration. For the low copy number, we used total 900 ng of DNA, with a transposase to transposon ratio of 1:2 and pRM105 to pRM106 ratio of 1:1. The day before integration, 500,000 hIPSCs were seeded in a 12 well-plate. On the day of integration, 300 ng of pRM105 and pRM106 each were mixed with 150 ng of transposase plasmids in separate Eppendorf tubes. For the high copy number integration, we used 5000 ng total DNA, with a transposase to transposon ratio of 1:3 and pRM105 to pRM106 ratio of 5:1. The day before integration, 300,000 hiPSCs were seeded in a 6-well plate. On the day of integration, 3125 ng pRM105 was mixed with 1042 ng transposase in an Eppendorf tube. In a separate tube, 625 ng of pRM106 was mixed with 208 ng of transposase. The two DNA mixtures were diluted to 25 *µ*L in Opti-MEM. Separate mixtures of 3 *µ*L Lipofectamine™ Stem Transfection Reagent (Invitrogen) in 47 *µ*L OPtiMEM (Gibco) were prepared per reaction. 25 *µ*L of LipoSTEM mixture was added to individual diluted DNA tubes. After 10 minutes of LipoSTEM incubation at room temperature, the two DNA mixtures were added to seeded cells drop-wise, one after another, allowing uncorrelated integration of the two plasmid components into hiPSCs. Instead of the transposase plasmid, nuclease-free water was used as a control to track and confirm fluorescence loss from transient transfection.

##### Sorting

Integrated cells were passaged for at least 14 days to confirm the stable integration of plasmid components into the hiPSC genome. Integrated cells were resuspended as single cells for sorting with CloneR™2 (StemCell Technologies). SONY MA900 cell sorter equipped with 405nm, 488nm, 561nm, and 638nm lasers was used to sort the desired cell populations in normal sort mode. Sorted polyclonal cell populations were collected in 5 ml tubes in mTeSR plus medium supplemented with CloneR™2 and Penicillin-Streptomycin (Sigma-Aldrich). In the first sort, cells positive for both EBFP2 and iRFP720 were collected, expanded, and were co-cultured with HEK sender cells to verify synNotch functionality. The cells were sorted for a second round to screen and select the desired populations (as explained in 2.4). The second round of sorting for was repeated two times.

### 4.4 Time-lapse Imaging of CHO cell co-culture for synNotch activation

CHO sender, receiver, and WT cells were detached from the culture dish using Versene (Gibco) and maintained at a concentration of 90,000 cells/mL. 50% receivers, 10% senders, and 40% WT cells (as fillers) were seeded in a 24-well plate at total volume of 500 *µ*L per well. Time-lapse images were acquired from 69 to 117 hours post incubation at every two-hour time interval using a Leica DM16000 Confocal Laser Scanning Microscope, in an incubation chamber. Images were captured with 10x objective. Bright-field and EBFP2 were imaged with 405 nm laser. 488 nm and 543 nm lasers were used to capture YFP and mKate fluorescence, respectively.

### 4.5 Single-cell analysis of synNotch activation dynamics

#### Cell Profiler Tracking

We quantified the sender-receiver interactions and synNotch activation of 359 individual receiver cells over a period of 48 hours with a pixel-based image analysis approach. First, we split up the composite fluorescent microscopy images into the 3 fluorescence channels: blue, yellow, and red. Blue fluorescence is expressed by all receivers, yellow fluorescence is expressed by senders, and red fluorescence is expressed by activated receivers. We used Cell Profiler’s `IdentifyPrimaryObjects`, `TrackObjects`, and `OverlayOutlines` modules to track the receiver and sender cells from the blue and yellow channels, respectively.

For the receiver cells, we used the following hyperparameters for the cell Profiler modules: typical object diameter 7-40 pixels, discard objects outside the diameter range, discard objects touching the border, global Otsu thresholding with two classes, threshold smoothing scale of 1.3488, threshold correction factor of 1.0, threshold lower and upper bounds of 0.3 and 1.0 respectively, log transform before thresholding, distinguish clumped objects based on intensity, draw dividing lines between clumped objects based on intensity, automatically calculate the size of smoothing filer for declumping, automatically calculate minimum allowed distance between local maxima, do not speed up by using lower-resolution image to find local maxima, do not display accepted local maxima, fill holes in identified objects after both thresholding and declumping, continue handling objects if excessive number of objects is identified, maximum pixel distance of 15 between objects in subsequent images to consider matches, color only display option, save color-coded image, and display outlines on a blank image. We passed the the time-series blue channel receiver images through these modules to produce time-series data for individual receiver cells that included their positions, and the pixels they occupied within each image.

For the sender cells, we used the following hyperparameters for the Cell Profiler modules: typical object diameter 7-50 pixels, discard objects outside the diameter range, do not discard objects touching the border, global Otsu thresholding with two classes, threshold smoothing scale of 1.3488, threshold correction factor of 1.0, threshold lower and upper bounds of 0.3 and 1.0 respectively, log transform before thresholding, distinguish clumped objects based on intensity, draw dividing lines between clumped objects based on intensity, use a smoothing filter of size 6 for declumping, automatically calculate minimum allowed distance between local maxima, do not speed up by using lower-resolution image to find local maxima, do not display accepted local maxima, fill holes in identified objects after both thresholding and declumping, continue handling objects if excessive number of objects is identified, maximum pixel distance of 15 between objects in subsequent images to consider matches, color only display option, save color-coded image, and display outlines on a blank image. We passed the time-series yellow channel sender images through these modules to produce time-series data showing the pixels that define the inner outline of the sender cells.

#### Quantifying Fluorescence and Interaction

We used the Cell Profiler tracking data to build two preliminary datasets, one for activated receivers and another for inactivated receivers, of time-series fluorescence and sender interaction data for individual receiver cells. We only consider receivers that were properly tracked by Cell Profiler for the entirety of the 48 hours. We determined by eye if a cell was properly tracked based on graphs similar to **Fig. 2c** that display a zoomed-in image of the cell at every timepoint. We calculated the fluorescence of a receiver at a given timepoint as the average red RGB value (on a scale from 0 to 1) of all pixels in the red channel that were occupied by the receiver cell at that time. We saved a receiver in the activated dataset if the raw mean fluorescence was greater than 0.03 at 48 hours, and less than 0.03 at hour zero to select receivers that started inactivated and became activated over the 48 hours. We saved a receiver in the inactivated dataset if the raw mean fluorescence was less than or equal to 0.03 at 48 hours, and less than 0.03 at hour zero to select receivers that started inactivated and remained inactivated over the 48 hours. We chose a threshold of 0.03 after determining that Cell Profiler is able to accurately differentiate the receivers from the background in the red fluorescence channel images when the threshold lower bound for `IdentifyPrimryObjects` is set to 0.03. In all graphs displaying mean red fluorescence of receivers (**Fig. 2c, Fig. 2d, Fig. 2e, Fig. 2f, Fig. 5e, Fig. SI2, Fig. SI3, Fig. SI4, Fig. SI6a, Fig. SI6c**), we normalize the fluorescence values by dividing by the maximum mean red fluorescence observed in a receiver at any time during the experiment.

To quantify the sender interaction the receivers experienced at each time, we consider the size and separation distance of the physical interfaces between sender and receiver membranes. Specifically, we took each pixel along the outline of a receiver cell and determined the euclidean distance to the nearest pixel on a sender outline. These values were passed through an exponential decay function and summed to determine the interaction level. Therefore, our formula for interaction at a given time is as follows:

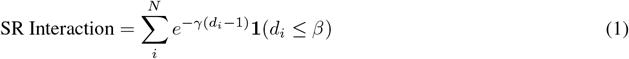

where *N* is the number of pixels on the outline of a receiver cell, *γ* is a distance-based signaling decay rate hyperparameter which we set to 1*/*3, *d*_*i*_ is the euclidean distance (in pixels) of the *i*th pixel on the receiver cell outline to the nearest pixel on the outline of a sender cell, **1** is an indicator function that is 1 when *d*_*i*_ ≤ *β* and 0 otherwise, and *β* is an interaction distance cutoff in pixels. We subtract 1 from *d*_*i*_ so that adjacent pixels (which have a euclidean distance of 1) are given a separation distance of 0. We set *β* = 10 and choose a signaling decay rate of *γ* = 1*/*3 so that sender outline pixels at a distance greater than 10 pixels (*≈* 15.2 µm) cannot contribute to the interaction sum. To normalize interaction values across experiments with different magnification than that of the original 48 hour experiment (**Fig. 2**), we first calculate the ratio of the new magnification to the magnification in the original experiment: 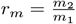. Here, *m*_1_ and *m*_2_ are the µm per pixel unit ratios in the fluorescence microscopy images of the original and new experiments respectively. Then, we use *r*_*m*_ in a modified version of (1) to calculate the cumulative interaction at a given time:

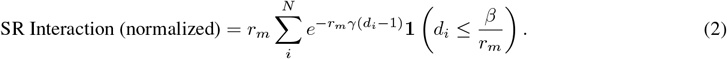

In this way, the chosen parameters (*γ* = 1*/*3, *β* = 10) and magnification of the original experiment are the baselines that we normalize to. We use this approach to calculate the interaction values in the experiment with micro-robot guided SynNotch activation **Fig. 5e**.

#### Activation Time Delay

While fluorescence threshold is a reasonable metric for determining activation, it fails to consider receivers that become activated towards the end of the 48 hours or those that slowly increase in mKate fluorescence after activation. In both of these cases, the final fluorescence may not exceed a given threshold or be visually apparent. However, as shown in **Fig. SI3**, it is possible to observe a trend of increasing fluorescence post-interaction for receivers that did not exceed the fluorescence threshold. Therefore, we use curve-fitting to make a final determination on which receivers are activated, inactivated, or removed from the dataset. Specifically, we fit a ReLU-like function to the fluorescence curves:

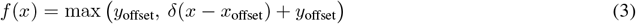

and determine the time of activation onset as the time at which *x* = *x*_offset_ where the slope of the function changes from zero to a positive number. We classified a receiver as activated if the following conditions were all met: the receiver experienced sender interaction before activation onset, the final mKate fluorescence was greater than the initial, and the receiver’s *x*_offset_ was greater than 0 and at least 3 timepoints before the end of the experiment (≤ 42 hours). We chose 42 hours as the cutoff to have at least 3 timesteps for which we could observe the fluorescence derivative. We classified a receiver as inactivated if any of the following conditions were met: the receiver did not experience sender interaction, the final mKate fluorescence was less than the initial, or the receiver’s *x*_offset_ was less than 3 timepoints before the end of the experiment (*>* 42 hours).

Finally, we sorted the activated receivers from least to most interaction, and the inactivated receivers from most to least interaction and added them into the finalized activated and inactivated receiver datasets in that order. To avoid bias, we do not add receivers to a dataset if they share a parent cell with another receiver already in that dataset. This means that activated receivers with low interaction will have priority over those with high interaction, and inactivated receivers with high interaction will have priority over those with low interaction. We choose these priorities to better understand the sender interaction threshold for receiver activation. With this approach, we produced a dataset of 59 activated receivers, and 300 inactivated receivers. We then use SciPy’s kernel-density estimate function on the interactions at each timepoint up to and including activation onset for each receiver, as shown by the orange curve in **Fig. SI2** and **Fig. SI3**. We calculate the activation delay time for each receivers by subtracting the activation onset time from the time of maximum KDE value (highest density/probability of interaction). Finally, we average the 59 activated receiver delay times (discarding 4 outliers greater than 24 hours) to obtain the average activation delay time of 6.7 *±* 4.6 hours.

### 4.6 Microrobots fabrication

We fabricated microrobots (MRs) using physical vapor deposition (PVD) technique. Using a custom made instrument from PVD Products (available at the University of Delaware’s Nanofabrication Facility), we spin-coated 4.8 *µ*m silica microspheres (CliniSciences, #SSD50003) onto a glass slide to form a monolayer of microspheres. Then, the microspheres were coated with a nickel layer using electron-beam physical vapor deposition to generate magnetic responsive MRs. We tried both 50 nm and 100 nm nickel layer thicknesses and used 100 nm nickel layer MRs for our study as we found it to be adequate to guide sender cells using our magnetic setup.

### 4.7 Sender cellbot preparation

We seeded CHO or HEK sender cells (10^5^ cells/dish) in a 35 mm Petri dish (MatTek Corporation, #NC9689997) and cultured them overnight. MRs were scratched off from the fabricated glass-slide and sonicated to make a well-dispersed solution. We pelleted MRs using a magnet and resuspended them in fresh 2 mL media before adding to the seeded cells. MR-cell coincubation was stored in an incubation chamber for 24 hours. MR-internalized cells were detached from the Petri dish and centrifuged to obtain the sender cellbots.

### 4.8 Cytocompatibility of Microrobots in CHO cells

To evaluate effects of MRs on sender cell morphology, we incubated MRs with seeded CHO cells for 24 hours. We imaged the MR-cell incubation using a ZOE fluorescent cell imager (Bio-Rad). and compared the images with a control cell culture without MRs. Cell viability after MR incubation was evaluated using flow cytometry (BD FACSAria Fusion). For flow analysis, we collected both supernatant and trypsinized MR-treated cells. After centrifugation, cells were resuspended in PBS and stained with 2% propidium iodide solution (Molecular probes, #P3566) and incubated for 5 minutes at room temperature before cell viability analysis. We used cells without MR incubation as positive control. To assess cell proliferation after MR incubation, we seeded sender cellbots and the control sender cell as single cells in a 96-well plate. We recorded cell division and microcolony formation by imaging the cells for 3 days using the ZOE fluorescence cell imager.

We evaluated the effect of magnetic actuation on sender cells and sender cellbots. Cells were exposed to magnetic field and actuated at 10 Hz for 15 minutes. Actuated cells were incubated for 24 hours. Microscopic images after actuation and 24 hours post incubation were acquired to monitor cell morphology. Cells without MR treatment and magnetic actuation were used as a positive control. We quantified cell viability after magnetic actuation using flow cytometry following 2% propidium iodide staining as described above.

### 4.9 Hardware in Magnetic controller system and Experimental set up

The hardware and experimental setup used in this study consists of a system that controls microrobots in open and closed-loop modes. For our implementation, the microrobots were actuated using a 3-pair orthogonal Helmholtz coil system, mounted on a Zeiss Axiovert 200M inverted microscope (**Fig 1b**). We designed and 3D printed the coil system, as explained in [39], where the small and medium coils have around 360 turns of 24 AWG copper wire, and the large coils have around 260 turns. The coil system is connected to an Arduino Mega 2560 which controls the current that goes to the coils. To compute the currents, we first use an acquisition system to visualize and analyze the working space, and then we extract the real-time feedback information. The acquisition system consists of a FLIR BFS-U3-50S5C-C camera and an algorithm to analyze each video frame implemented in a Jetson Xavier NX. Finally, the real-time feedback information is gathered from the visualized environment and sent to the Arduino Mega which outputs the signal. This system works as it is for the closed-loop control mode and skips the real-time feedback information when operating in open-loop control mode. In this specific scenario, the currents are determined by the movement of a wireless X-box controller. However, both modes use the camera, as the visualization on the microscopic scale is done through the camera. The speed of a microrobot or cellbot was estimated in real time using custom-made tracking and detection software. The software calculates the distance a microrobot or cellbot travels in 15 frames, and converts the distance in pixels to a distance in micrometers using microscope camera specifications. This distance is then divided by the time it took to capture 15 frames. This results in an average speed the microrobot or cellbot travels over certain number of frames.

### 4.10 CHO sender cellbot co-culture with receivers for synNotch activation

CHO sender cellbots were co-cultured with the receiver cells at 1:1 ratio in a 35 mm Petri dish. We used CHO senders and WT CHO cells as positive and negative controls, respectively. The cells were imaged to validate synNotch activation after 48 hours with a confocal laser scanning microscope (Zeiss LSM880) using 40X oil immersion objective. We used 405 nm, 514 nm, and 594 nm lasers to detect EBFP2, YFP, and mKate, respectively.

We quantified synNotch activation of sender-receiver co-culture using flow cytometry. Sender cellbots, senders and WT cells were co-cultured with the receivers at 1:1 ratio in a 12-well plate at a total cell density of 400,000 cells per well. After 48 hours, we detached the cells from the culture dish using Versene and resuspended them in PBS supplemented with 2% FBS for flow cytometry. Flow cytometry data was collected on a BD LSR Fortessa located in the MIT Synthetic Biology Center. FACSDiva software (BD) was used for the initial collection of 60,000–80,000 events in tube mode. For the acquisition, a population containing single live cells was determined based on forward and side scatters. We detected the expression of fluorescent proteins using the following settings: 405 nm laser with 450/50 nm filter (‘Pacific Blue’) for measuring EBFP2, 488 laser with 530/30 nm filter (‘FITC’) for measuring YFP and 561 nm laser with 610/20 nm filter (‘PE-Texas-Red’) for measuring mKate. We analyzed the flow cytometry data using FlowJo version 10.0. Briefly, the population of live cells was determined by plotting single cell events on forward and side scatter. Cross channel fluorescent bleed through was corrected using single-color controls established by transfecting fluorescent proteins expressed by relevant constitutive promoters. Graphs were generated and all statistical analyses were performed using GraphPad Prism 10 (GraphPad Software).

### 4.11 MR-guided synNotch activation in CHO receiver cells

CHO Sender cellbots were prepared as described previously. Receiver cells were cultured in a 35 mm petri dish up to 40-50% confluency. Receiver cells were replenished with fresh media and were placed on the magnetic control platform (Fig 3c). Approximately 50 sender cellbots were added to the receiver cell culture and guided to desired locations at a frequency between 5-10 Hz. Cells were left undisturbed for 1 hour to allow the sender cellbots to adhere to the petri dish. Time-lapse fluorescence images were acquired for 26 hours at every two-hour time interval, using a Leica DM16000 Confocal Laser Scanning Microscope, in an incubation chamber. We acquired images using a 10X objective using 405 nm, 488 nm and 543 nm lasers to detect EBFP2, YFP and mKate, respectively.

### 4.12 Transient poly-transfection of synNotch receiver circuits in HEK cell

SynNotch-based ETV2 expressing circuits were characterized using poly-transfection based on a previously published protocol [43] using Lipofectamine™ 3000 Transfection Reagent (Invitrogen). Briefly, plasmids, pRM105 and pM106, were maintained at concentrations of 100 ng/*µ*L. 450 ng of each plasmid was diluted in OptiMEM (0.05 *µ*L/ng of DNA) and mixed with P3000 (0.0022 *µ*L/ng of DNA) in separate Eppendorf tubes. Lipofectamine™ 3000 (0.0022 *µ*L/ng of DNA) was diluted in OptiMEM (0.05 *µ*L/ng of DNA) separately and added to the DNA P3000 mixtures. The two transfection mixes were then added dropwise to freshly seeded 150,000 HEK293FT cells in a 24-well plate, one after the other, to allow uncorrelated plasmid transfection. After 24 hours of transfection, resulting receiver cells were detached using Versene and co-cultured with stable HEK senders (or WT HEK cells as a negative control) at 1:1 ratio in a 24-well plate for synNotch activation. After 48 hours of co-culture, cells were detached and collected for flow cytometry using a BD LSR Fortessa as described in section 4.10.

Using FACSDiva software (BD), we determined single live cells based on initial forward and side scatters. We collected approximately 200,000 events in tube mode. For the detection of fluorescent protein expression, we used the following settings: 405 nm laser with 450/50 nm filter (‘Pacific Blue’) for measuring EBFP2, 488 nm laser with 530/30 filter (‘FITC’) for measuring YFP, 561 nm laser with 610/20 nm filter (‘PE-Texas-Red’) for measuring mKO2 and 637 nm laser with 710/50 nm filter (‘Alexa Fluor 700’) for measuring iRFP720. We analyzed the flow cytometry data using Cytoflow-1.2. Briefly, using wild-type HEK cells, we set the morphological gating to isolate the single-cell events based on forward and side scatters. Cross channel fluorescent bleed through was corrected using single- and multi-color controls, which were prepared by transfecting HEK cells with fluorescent proteins expressed by relevant constitutive promoters in the poly-transfection ratios. Binning was performed on iRFP720 and EBFP2 as an indication of plasmid copy numbers in transfected cells. In each bin, the mean mKO2 fluorescent intensity was estimated to quantify synNotch activation for specific plasmid copy numbers. Fold-change in mKO2 expression between the sample and negative control was calculated for individual bins. Graphs were generated using GraphPad Prism 10 (GraphPad Software).

### 4.13 HEK sender and hiPSC receiver co-culture for synNotch activation

For synNotch activation-based ETV2 expression in hiPSC receivers, we co-cultured the selected hiPSC receive cell line with HEK senders. HiPSC receivers were seeded on matrigel-coated 24-well plates at a cell density of 100,000 cells per well. After 24 hours, 25,000 HEK sender cells were added per well in mTeSR media to maintain the co-culture. WT HEK cells were co-cultured with hiPSC receivers as a negative control. Two days after the co-culture, we changed the media in the wells before acquiring fluorescence images using a Leica DM16000 confocal laser scanning microscope with a 40X oil immersion objective. We used 405 nm, 488 nm, 543 nm, and 633 nm lasers to acquire fluorescence from EBFP2, YFP, mKO2, and iRFP720 expression, respectively.

### 4.14 Immunofluorescence to verify EC-specific markers

hiPSC receiver cells were seeded in a Matrigel-coated 24-well plate at a density of 100,000 cells/well. After 24 hours, 25,000 HEK sender cells were co-cultured with the receiver cells. The co-culture was maintained in mTeSR media for eight days. WT HEK cells, instead of HEK senders, were co-cultured with hiPSC receivers as a negative control. After eight days, the cells were fixed by incubating with 100 ul 4% paraformaldehyde (Electron Microscopy Sciences) for 10 min at room temperature. The cells were then permeabilized using Triton X-100 (Sigma-Aldrich) for 10 minutes at room temperature. Next, the wells containing permeabilized cells were blocked using 1% Bovine Serum Albumin (New England Biolabs) supplemented with 0.3M glycine (Fisher BioReagents™) for two hours at room temperature. The blocked wells were then washed and incubated with Anti-VE-Cadherin primary Ab (Abcam, #313632) at 4°C overnight. The cells were then washed and incubated with donkey anti-rabbit IgG (H+L) Highly Cross-Adsorbed secondary antibody, Alexa Fluor™ 488 (Invitrogen, #A-21206) for one hour at room temperature in the dark. Cells were washed to remove unbound antibodies before imaging. Fluorescence images of the antibody stained samples were acquired using a Leica DM16000 confocal laser scanning microscope with a 40X oil immersion objective. We used 405 nm, 476 nm, 488 nm, 543 nm, and 633 nm lasers to acquire fluorescence of eBFP2, AF488, YFP, mKO2, and iRFP720, respectively.

### 4.15 RT-qPCR to verify hiPSC differentiation

RNA was extracted using Monarch® Total RNA Miniprep Kit (NEB). 200 ng of RNA was reverse transcribed using iScript™ gDNA Clear cDNA Synthesis Kit (Bio-Rad). RT-qPCR reactions using the resulting cDNA were prepared using FastSYBR mixture (CoWin Biosciences). RT-qPCR was performed using a LightCycler 96 (Roche). Data analysis was done using the LightCycler 96 software and Microsoft Excel. The relative amounts of each gene were normalized to that of the expression of the housekeeping gene, GAPDH. A complete list of primers used can be found in **Table SI3**.

### 4.16 Microrobot-guided hiPSC differentiation

Half of 35 mm petri dishes were coated with Matrigel, which we used to seed hiPSC receivers. After overnight incubation, the receiver cell culture was moved to the magnetic control platform. HEK sender cellbots were prepared as described above in section 4.7. Approximately ten thousand sender cellbots were added to the non-matrigel coated side of the dish and guided to the target location of the receivers at a frequency of 5 Hz. After the sender cellbot guidance, the culture was moved carefully to a nearby cell culture incubation chamber to avoid untargeted movement of sender cellbots. Fluorescence images of the co-culture were acquired at several time points using a Leica DM16000 confocal laser scanning microscope with both 10X and 40X objectives. After eight days of co-culture, the cells were fixed for immunofluorescence study to validate hiPSC differentiation into ECs.

## 5 Acknowledgements

This work is supported by Growing Convergence Research at NSF. We thank the Koch Institute Flow Cytometry Core Facility at MIT and acknowledge D. Ray and C. Haase-Pettingell for administrative support. The schematics in the study were prepared using BioRender.

## Notes

### Competing Interest Statement

The authors have declared no competing interest.

## References

[1] Christian Mandrycky, Zongjie Wang, Keekyoung Kim, and Deok-Ho Kim. 3d bioprinting for engineering complex tissues. Biotechnology advances, 34(4):422–434, 2016.

[2] Xiaolei Yin, Benjamin E Mead, Helia Safaee, Robert Langer, Jeffrey M Karp, and Oren Levy. Engineering stem cell organoids. Cell stem cell, 18(1):25–38, 2016.

[3] Jonathan A Brassard and Matthias P Lutolf. Engineering stem cell self-organization to build better organoids. Cell stem cell, 24(6):860–876, 2019.

[4] Akinao Nose, Akira Nagafuchi, and Masatoshi Takeichi. Expressed recombinant cadherins mediate cell sorting in model systems. Cell, 54(7):993–1001, 1988.

[5] Duke Duguay, Ramsey A Foty, and Malcolm S Steinberg. Cadherin-mediated cell adhesion and tissue segregation: qualitative and quantitative determinants. Developmental biology, 253(2):309–323, 2003.

[6] Ramsey A Foty and Malcolm S Steinberg. The differential adhesion hypothesis: a direct evaluation. Developmental biology, 278(1):255–263, 2005.

[7] Elise Cachat, Weijia Liu, Kim C Martin, Xiaofei Yuan, Huabing Yin, Peter Hohenstein, and Jamie A Davies. 2-and 3-dimensional synthetic large-scale de novo patterning by mammalian cells through phase separation. Scientific reports, 6(1):20664, 2016.

[8] Elise Cachat, Weijia Liu, and Jamie A Davies. Synthetic self-patterning and morphogenesis in mammalian cells: a proof-of-concept step towards synthetic tissue development. Engineering Biology, 1(2):71–76, 2017.

[9] Fokion Glykofrydis, Elise Cachat, Ieva Berzanskyte, Elaine Dzierzak, and Jamie A Davies. Bioengineering self-organizing signaling centers to control embryoid body pattern elaboration. ACS Synthetic Biology, 10(6):1465–1480, 2021.

[10] Jesse Tordoff, Matej Krajnc, Nicholas Walczak, Matthew Lima, Jacob Beal, Stanislav Shvartsman, and Ron Weiss. Incomplete cell sorting creates engineerable structures with long-term stability. Cell Reports Physical Science, 2(1), 2021.

[11] Noreen Wauford, Akshay Patel, Jesse Tordoff, Casper Enghuus, Andrew Jin, Jack Toppen, Melissa L Kemp, and Ron Weiss. Synthetic symmetry breaking and programmable multicellular structure formation. Cell Systems, 14(9):806–818, 2023.

[12] George Chao, Timothy M Wannier, Clair Gutierrez, Nathaniel C Borders, Evan Appleton, Anjali Chadha, Tina Lebar, and George M Church. helixcam: A platform for programmable cellular assembly in bacteria and human cells. Cell, 185(19):3551–3567, 2022.

[13] Adam J Stevens, Andrew R Harris, Josiah Gerdts, Ki H Kim, Coralie Trentesaux, Jonathan T Ramirez, Wesley L McKeithan, Faranak Fattahi, Ophir D Klein, Daniel A Fletcher, et al. Programming multicellular assembly with synthetic cell adhesion molecules. Nature, 614(7946):144–152, 2023.

[14] Evan Appleton, Noushin Mehdipour, Tristan Daifuku, Demarcus Briers, Iman Haghighi, Michaël Moret, George Chao, Timothy Wannier, Anush Chiappino-Pepe, Jeremy Huang, et al. Algorithms for autonomous formation of multicellular shapes from single cells. ACS Synthetic Biology, 2024.

[15] Pulin Li, Joseph S Markson, Sheng Wang, Siheng Chen, Vipul Vachharajani, and Michael B Elowitz. Morphogen gradient reconstitution reveals hedgehog pathway design principles. Science, 360(6388):543–548, 2018.

[16] Satoshi Toda, Wesley L McKeithan, Teemu J Hakkinen, Pilar Lopez, Ophir D Klein, and Wendell A Lim. Engineering synthetic morphogen systems that can program multicellular patterning. Science, 370(6514):327–331, 2020.

[17] Kristina S Stapornwongkul, Marc de Gennes, Luca Cocconi, Guillaume Salbreux, and Jean-Paul Vincent. Patterning and growth control in vivo by an engineered gfp gradient. Science, 370(6514):321–327, 2020.

[18] Gustav Y Cederquist, James J Asciolla, Jason Tchieu, Ryan M Walsh, Daniela Cornacchia, Marilyn D Resh, and Lorenz Studer. Specification of positional identity in forebrain organoids. Nature biotechnology, 37(4):436–444, 2019.

[19] Ryoji Sekine, Tatsuo Shibata, and Miki Ebisuya. Synthetic mammalian pattern formation driven by differential diffusivity of nodal and lefty. Nature communications, 9(1):5456, 2018.

[20] Mitsuhiro Matsuda, Makito Koga, Knut Woltjen, Eisuke Nishida, and Miki Ebisuya. Synthetic lateral inhibition governs cell-type bifurcation with robust ratios. Nature communications, 6(1):6195, 2015.

[21] Mitsuhiro Matsuda, Makito Koga, Eisuke Nishida, and Miki Ebisuya. Synthetic signal propagation through direct cell-cell interaction. Science signaling, 5(220):ra31–ra31, 2012.

[22] Leonardo Morsut, Kole T Roybal, Xin Xiong, Russell M Gordley, Scott M Coyle, Matthew Thomson, and Wendell A Lim. Engineering customized cell sensing and response behaviors using synthetic notch receptors. Cell, 164(4):780–791, 2016.

[23] Marco Santorelli, Pranav S Bhamidipati, Andriu Kavanagh, Victoria A MacKrell, Trusha Sondkar, Matt Thomson, and Leonardo Morsut. Control of spatio-temporal patterning via cell density in a multicellular synthetic gene circuit. bioRxiv, pages 2022–10, 2022.

[24] Satoshi Toda, Lucas R Blauch, Sindy KY Tang, Leonardo Morsut, and Wendell A Lim. Programming self-organizing multicellular structures with synthetic cell-cell signaling. Science, 361(6398):156–162, 2018.

[25] Mattias Malaguti, Rosa Portero Migueles, Jennifer Annoh, Daina Sadurska, Guillaume Blin, and Sally Lowell. Synpl: Synthetic notch pluripotent cell lines to monitor and manipulate cell interactions in vitro and in vivo. Development, 149(12):dev200226, 2022.

[26] Mher Garibyan, Tyler Hoffman, Thijs Makaske, Stephanie K Do, Yifan Wu, Brian A Williams, Alexander R March, Nathan Cho, Nicolas Pedroncelli, Ricardo Espinosa Lima, et al. Engineering programmable material-to-cell pathways via synthetic notch receptors to spatially control differentiation in multicellular constructs. Nature Communications, 15(1):5891, 2024.

[27] Mark A Skylar-Scott, Jeremy Y Huang, Aric Lu, Alex HM Ng, Tomoya Duenki, Songlei Liu, Lucy L Nam, Sarita Damaraju, George M Church, and Jennifer A Lewis. Orthogonally induced differentiation of stem cells for the programmatic patterning of vascularized organoids and bioprinted tissues. Nature biomedical engineering, 6(4):449–462, 2022.

[28] Heath E Johnson, Yogesh Goyal, Nicole L Pannucci, Trudi Schüpbach, Stanislav Y Shvartsman, and Jared E Toettcher. The spatiotemporal limits of developmental erk signaling. Developmental cell, 40(2):185–192, 2017.

[29] Daniel Krueger, Emiliano Izquierdo, Ranjith Viswanathan, Jonas Hartmann, Cristina Pallares Cartes, and Stefano De Renzis. Principles and applications of optogenetics in developmental biology. Development, 146(20):dev175067, 2019.

[30] Jonas Hartmann, Daniel Krueger, and Stefano De Renzis. Using optogenetics to tackle systems-level questions of multicellular morphogenesis. Current opinion in cell biology, 66:19–27, 2020.

[31] Guillermo Martínez-Ara, Núria Taberner, Mami Takayama, Elissavet Sandaltzopoulou, Casandra E Villava, Miquel Bosch-Padrós, Nozomu Takata, Xavier Trepat, Mototsugu Eiraku, and Miki Ebisuya. Optogenetic control of apical constriction induces synthetic morphogenesis in mammalian tissues. Nature communications, 13(1):5400, 2022.

[32] Nicole A Repina, Hunter J Johnson, Xiaoping Bao, Joshua A Zimmermann, David A Joy, Shirley Z Bi, Ravi S Kane, and David V Schaffer. Optogenetic control of wnt signaling models cell-intrinsic embryogenic patterning using 2d human pluripotent stem cell culture. Development, 150(14), 2023.

[33] Ivano Legnini, Lisa Emmenegger, Alessandra Zappulo, Agnieszka Rybak-Wolf, Ricardo Wurmus, Anna Oliveras Martinez, Cledi Cerda Jara, Anastasiya Boltengagen, Talé Hessler, Guido Mastrobuoni, et al. Spatiotemporal, optogenetic control of gene expression in organoids. Nature Methods, 20(10):1544–1552, 2023.

[34] Deasung Jang, Jinwon Jeong, Hyeonseok Song, and Sang Kug Chung. Targeted drug delivery technology using untethered microrobots: A review. Journal of Micromechanics and Microengineering, 29(5):053002, 2019.

[35] Wenjun Chen, Hao Zhou, Bin Zhang, Qinghua Cao, Bo Wang, and Xing Ma. Recent progress of micro/nanorobots for cell delivery and manipulation. Advanced Functional Materials, 32(18):2110625, 2022.

[36] Lidong Yang and Li Zhang. Motion control in magnetic microrobotics: From individual and multiple robots to swarms. Annual Review of Control, Robotics, and Autonomous Systems, 4(1):509–534, 2021.

[37] David R Stirling, Madison J Swain-Bowden, Alice M Lucas, Anne E Carpenter, Beth A Cimini, and Allen Goodman. Cellprofiler 4: improvements in speed, utility and usability. BMC bioinformatics, 22:1–11, 2021.

[38] David Rivas, Sudipta Mallick, Max Sokolich, and Sambeeta Das. Cellular manipulation using rolling microrobots. In 2022 International Conference on Manipulation, Automation and Robotics at Small Scales (MARSS), pages 1–6. IEEE, 2022.

[39] Max Sokolich, David Rivas, Yanda Yang, Markos Duey, and Sambeeta Das. Modmag: A modular magnetic micro-robotic manipulation device. MethodsX, 10:102171, 2023.

[40] Chengwei Ye, Jia Liu, Xinyu Wu, Ben Wang, Li Zhang, Yuanyi Zheng, and Tiantian Xu. Hydrophobicity influence on swimming performance of magnetically driven miniature helical swimmers. Micromachines, 10(3):175, 2019.

[41] Rimpei Morita, Mayu Suzuki, Hidenori Kasahara, Nana Shimizu, Takashi Shichita, Takashi Sekiya, Akihiro Kimura, Ken-ichiro Sasaki, Hideo Yasukawa, and Akihiko Yoshimura. Ets transcription factor etv2 directly converts human fibroblasts into functional endothelial cells. Proceedings of the National Academy of Sciences, 112(1):160–165, 2015.

[42] Irina Elcheva, Vera Brok-Volchanskaya, Akhilesh Kumar, Patricia Liu, Jeong-Hee Lee, Lilian Tong, Maxim Vodyanik, Scott Swanson, Ron Stewart, Michael Kyba, et al. Direct induction of haematoendothelial programs in human pluripotent stem cells by transcriptional regulators. Nature communications, 5(1):4372, 2014.

[43] Jeremy J Gam, Breanna DiAndreth, Ross D Jones, Jin Huh, and Ron Weiss. A ‘poly-transfection’method for rapid, one-pot characterization and optimization of genetic systems. Nucleic Acids Research, 47(18):e106–e106, 2019.

[44] Katherine A Kiwimagi, Justin H Letendre, Benjamin H Weinberg, Junmin Wang, Mingzhe Chen, Leandro Watanabe, Chris J Myers, Jacob Beal, Wilson W Wong, and Ron Weiss. Quantitative characterization of recombinase-based digitizer circuits enables predictable amplification of biological signals. Communications biology, 4(1):875, 2021.

[45] Masakazu Inada, Genya Izawa, Wakako Kobayashi, and Masayuki Ozawa. 293 cells express both epithelial as well as mesenchymal cell adhesion molecules. International journal of molecular medicine, 37(6):1521–1527, 2016.

[46] Francesca Soncin and Christopher M Ward. The function of e-cadherin in stem cell pluripotency and self-renewal. Genes, 2(1):229–259, 2011.

[47] Hongyan Zhang, Tomoko Yamaguchi, Yasuhiro Kokubu, and Kenji Kawabata. Transient etv2 expression promotes the generation of mature endothelial cells from human pluripotent stem cells. Biological and Pharmaceutical Bulletin, 45(4):483–490, 2022.

[48] Xavier Duportet, Liliana Wroblewska, Patrick Guye, Yinqing Li, Justin Eyquem, Julianne Rieders, Tharathorn Rimchala, Gregory Batt, and Ron Weiss. A platform for rapid prototyping of synthetic gene networks in mammalian cells. Nucleic acids research, 42(21):13440–13451, 2014.

[49] Ernst Weber, Carola Engler, Ramona Gruetzner, Stefan Werner, and Sylvestre Marillonnet. A modular cloning system for standardized assembly of multigene constructs. PloS one, 6(2):e16765, 2011.

[50] Lei Wang, Wenlong Xu, Shun Zhang, Gregory C Gundberg, Christine R Zheng, Zhengpeng Wan, Kamila Mustafina, Fabio Caliendo, Hayden Sandt, Roger Kamm, et al. Sensing and guiding cell-state transitions by using genetically encoded endoribonuclease-mediated microrna sensors. Nature Biomedical Engineering, pages 1–14, 2024.

[51] Shuang Zhao, Enze Jiang, Shuangshuang Chen, Yuan Gu, Anna Junjie Shangguan, Tangfeng Lv, Liguo Luo, and Zhenghong Yu. Piggybac transposon vectors: the tools of the human gene encoding. Translational lung cancer research, 5(1):120, 2016.

