## Supplementary Information for "Magnetized Cellbots to Spatiotemporally Control Differentiation of Human-Induced Pluripotent Stem Cells"

March 25, 2025

#### 1 Rotating Magnetic Field Strategy

The following are equations for modeling the sinusoidal signals of a rotating magnetic field [1]:

$$\mathbf{B}_x = \cos(\gamma)\cos(\alpha)\cos(2\pi ft) + \sin(\alpha)\sin(2\pi ft), \tag{1}$$

$$\mathbf{B}_y = -\cos(\gamma)\sin(\alpha)\cos(2\pi ft) + \cos(\alpha)\sin(2\pi ft), \tag{2}$$

$$\mathbf{B}_z = \sin(\gamma)\cos(2\pi ft), \tag{3}$$

The azimuthal angle of the rotation axis is controlled by adjusting the variable $\alpha$ from $0 < \alpha < 360°$, which determines the direction of the magnetic microrobot. The polar angle of the rotation axis is controlled by adjusting the variable $\gamma$ from $0 < \gamma < 180°$, which controls the rolling actuation on planar xy surfaces $\gamma = 90°$. Finally, the frequency of the rotating magnetic field is controlled by adjusting the variable $f$ to modulate the speed of a MR or sender cellbot. $\gamma$, and $f$ were adjusted in the control software while $\alpha$ was adjusted using the left joystick of a PS4 gaming controller. Thus, the position of a MR or sender cellbot can be precisely controlled using the joystick of a gaming controller.

#### 2 Step Out Frequency

When the rotation frequency of the magnetic field is below the step-out frequency, the driving magnetic torque is large enough to counterbalance the resistive fluidic drag torque, which enables the object to rotate synchronously with the magnetic field. Its rolling velocity increases in an approximately linear pattern with respect to the increasing rotation frequency. When the rotation frequency of the magnetic field is beyond the step-out frequency, the magnetic torque is insufficient to offset the drag torque, therefore the roller rotates asynchronously with the applied field, and its rolling velocity undergoes a drastic and non-linear decline as the rotation frequency is increased. [2].

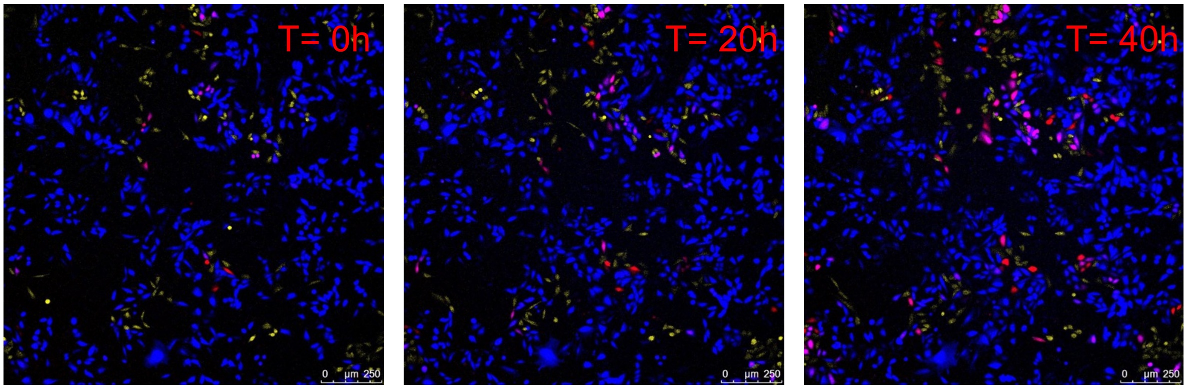

Figure SI1: Time-lapse images of synNotch activation in CHO sender and receiver cell co-culture.

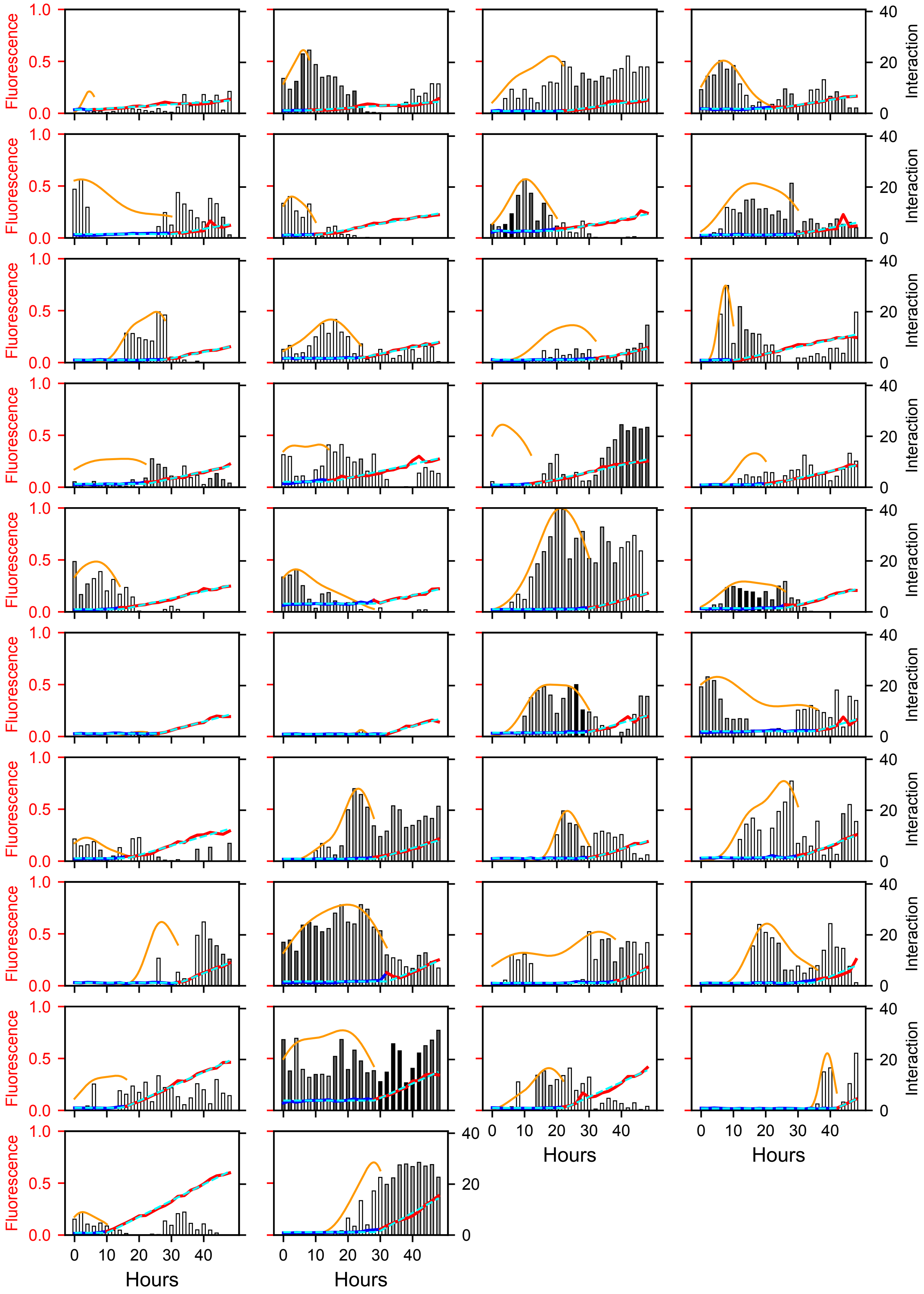
Fluorescence
Interaction
Hours
0.0
0.5
1.0
0
20
40
0 10 20 30 40

Figure SI2: **High Fluorescence Activated Receivers.** A set of graphs showing the interaction and normalized fluorescence values of 38 activated receiver cells over a period of 48 hours. These receivers have a raw fluorescence value greater than 0.03 (normalized value $\approx 0.1$) at 48 hours. The blue and red solid line is the cell's pixelwise mean red fluorescence at each timepoint, the indicator of mKate expression. This line is partitioned into blue and red sections corresponding to pre- and post- onset of activation respectively. Each bar in the graph represents the level of sender interaction with the receiver cell which is calculated based on the length of and distance between sender-receiver interfaces. Bars are grayscale-coded based on number of sender cells in close proximity to the receiver (white=1, light gray=2, dark gray=3, and black=4). The dashed cyan line in each plot is the ReLU function that was fit to the receiver's fluorescence data to determine the time of activation onset, and the derivative of fluorescence after activation. The orange curve is the gaussian kernel density estimate (KDE) of the receiver's probability density function for pre-activation interaction. This is used to find the timepoint with highest sender interaction density (highest likelihood of interaction). We normalize the KDE values to fall within the range of interaction in the plots.

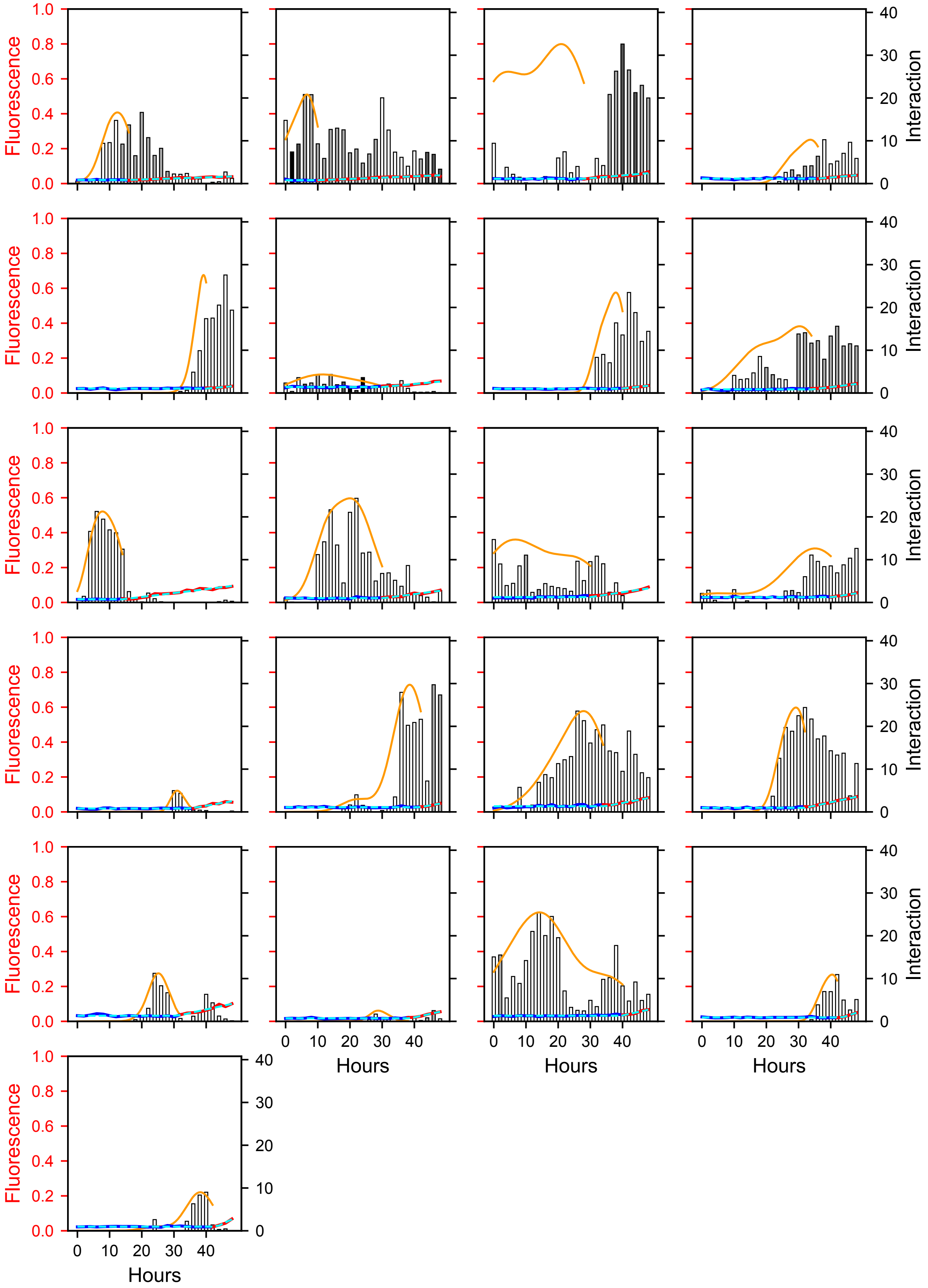
Fluorescence
Interaction
Hours
1.0
0.8
0.6
0.4
0.2
0.0
40
30
20
10
0
0 10 20 30 40

Figure SI3: **Low Fluorescence Activated Receivers.** A set of graphs showing the interaction and normalized fluorescence values of 21 activated receiver cells over a period of 48 hours. These receivers have a raw fluorescence value less than or equal to 0.03 (normalized value $\approx 0.1$) at 48 hours. The blue and red solid line is the cell's pixelwise mean red fluorescence at each timepoint, the indicator of mKate expression. This line is partitioned into blue and red sections corresponding to pre- and post- onset of activation respectively. Each bar in the graph represents the level of sender interaction with the receiver cell which is calculated based on the length of and distance between sender-receiver interfaces. Bars are grayscale-coded based on number of sender cells in close proximity to the receiver (white=1, light gray=2, dark gray=3, and black=4). The dashed cyan line in each plot is the ReLU function that was fit to the receiver's fluorescence data to determine the time of activation onset, and the derivative of fluorescence after activation. The orange curve is the gaussian kernel density estimate (KDE) of the receiver's probability density function for pre-activation interaction. This is used to find the timepoint with highest sender interaction density (highest likelihood of interaction). We normalize the KDE values to fall within the range of interaction in the plots.

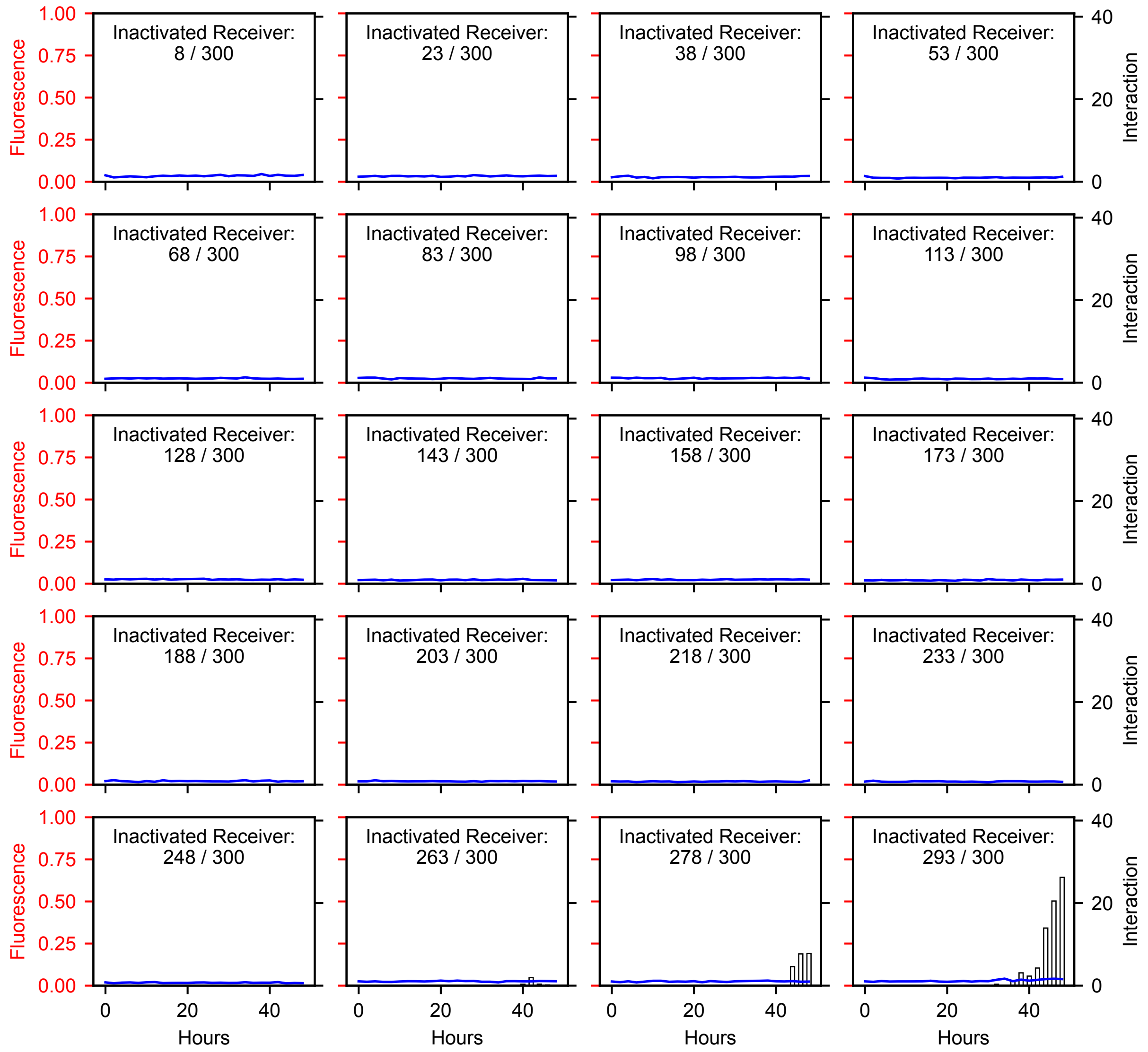


Figure SI4: **Inactivated Receivers.** A set of graphs showing the interaction and normalized fluorescence values of 20 representative inactivated receiver cells of the 300 total inactivated receivers. The receivers are sorted from least to most interaction, and the numbering indicates their sorted place in the set where 1 is the least interaction, and 300 is the most interaction. The blue solid line is the cell's pixelwise mean red fluorescence at each timepoint, the indicator of mKate expression. Each bar in the graph represents the level of sender interaction with the receiver cell which is calculated based on the length of and distance between sender-receiver interfaces. Bars are grayscale-coded based on number of sender cells in close proximity to the receiver (white=1, light gray=2, dark gray=3, and black=4).

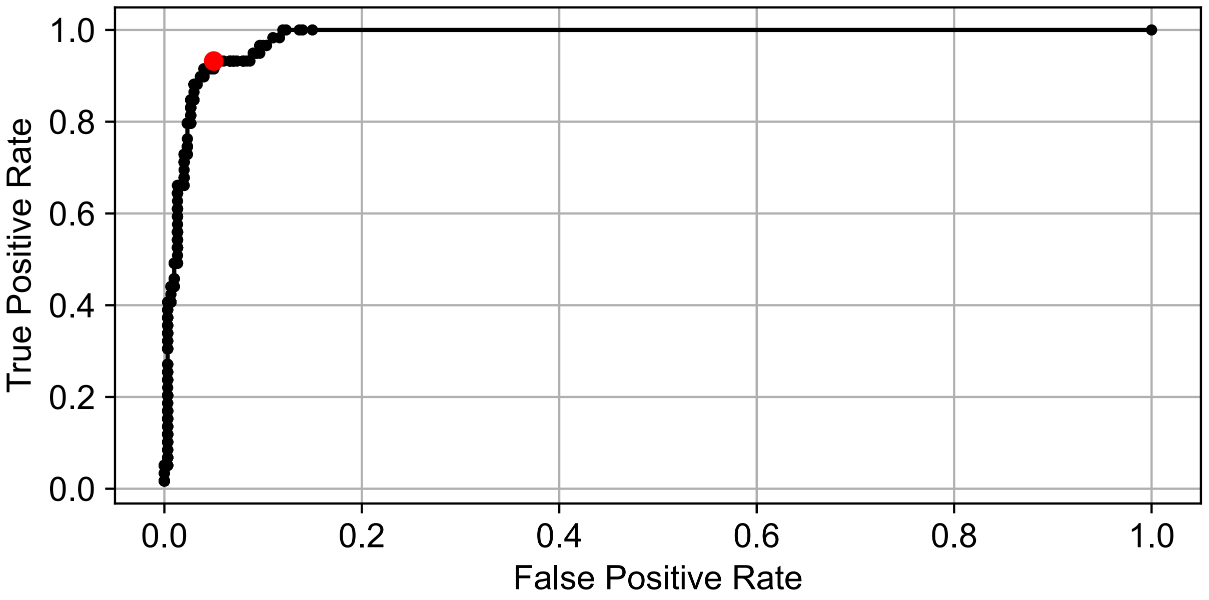


Figure SI5: A Receiver-Operating Characteristic (ROC) curve for predicting whether a receiver is activated or inactivated based on a cumulative interaction threshold. Cumulative interaction is calculated as the sum of interactions over 0 to 46 hours. Each data point represents the true positive and false positive rates of using a particular cumulative interaction threshold to classify receivers as activated or inactivated. If a receiver's cumulative interaction was greater than or equal to the threshold it was classified as activated, and otherwise was classified as inactivated. In this ROC curve, the true positives are the activated receivers with cumulative interaction greater than or equal to the threshold, the false positives are the inactivated receivers with cumulative interaction greater than or equal to the threshold, the true negatives are the inactivated receivers with cumulative interaction less than the threshold, and the false negatives are the activated receivers with cumulative interaction less than the threshold. To generate the thresholds for the ROC curve, we sampled 1000 evenly-spaced data points from 0 to the maximum cumulative interaction of any receiver (491.4). We calculated the euclidean distance of each point on the ROC curve to the perfect classifier ($[0, 1]$ on the plot) and determined that a threshold of 30 cumulative interaction corresponded to the point minimally distant from the perfect classifier (shown in red).

| | **Significant Interaction?** | |
|---|---|---|
| | ***Yes*** | ***No*** |
| **# Activated** | 55 | 4 |
| **# Inactivated** | 15 | 285 |

Table SI1: A table showing the numbers of activated and inactivated receivers with cumulative interaction (over 0–46 hours) above/below a threshold 30. We determine this threshold based on the ROC curve in **Fig. SI5**.

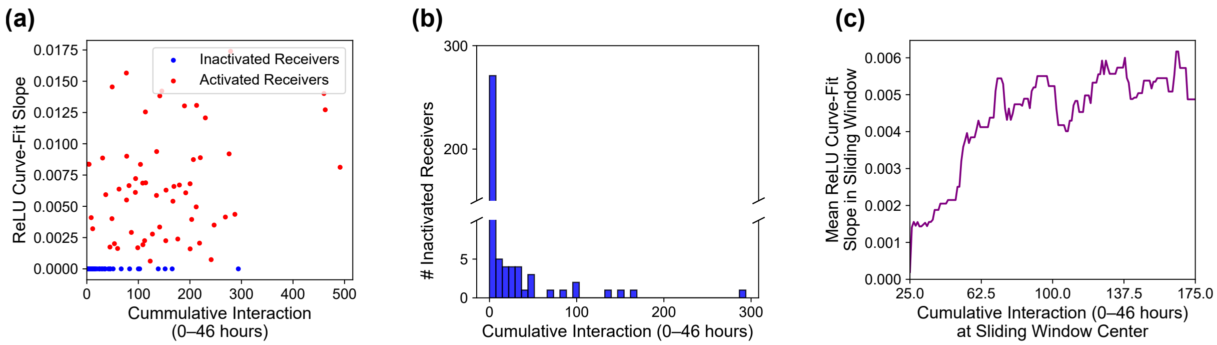


Figure SI6: (**a**) A scatter plot of the ReLU fluorescence curve-fit slope against cumulative interaction before activation. The slopes of all inactivated receiver cells will be 0, and their cumulative interaction is measured from 0 to 46 hours to account for each interaction timepoint that could have induced their activation. (**b**) A histograms of the number of inactivated receivers against from 0 to 46 hours. (**c**) A sliding-window plot showing the mean ReLU fluorescence curve-fit slope within the sliding window against the cumulative interaction at the center of the sliding window. A window size of 50 and a stride of 1 were used.

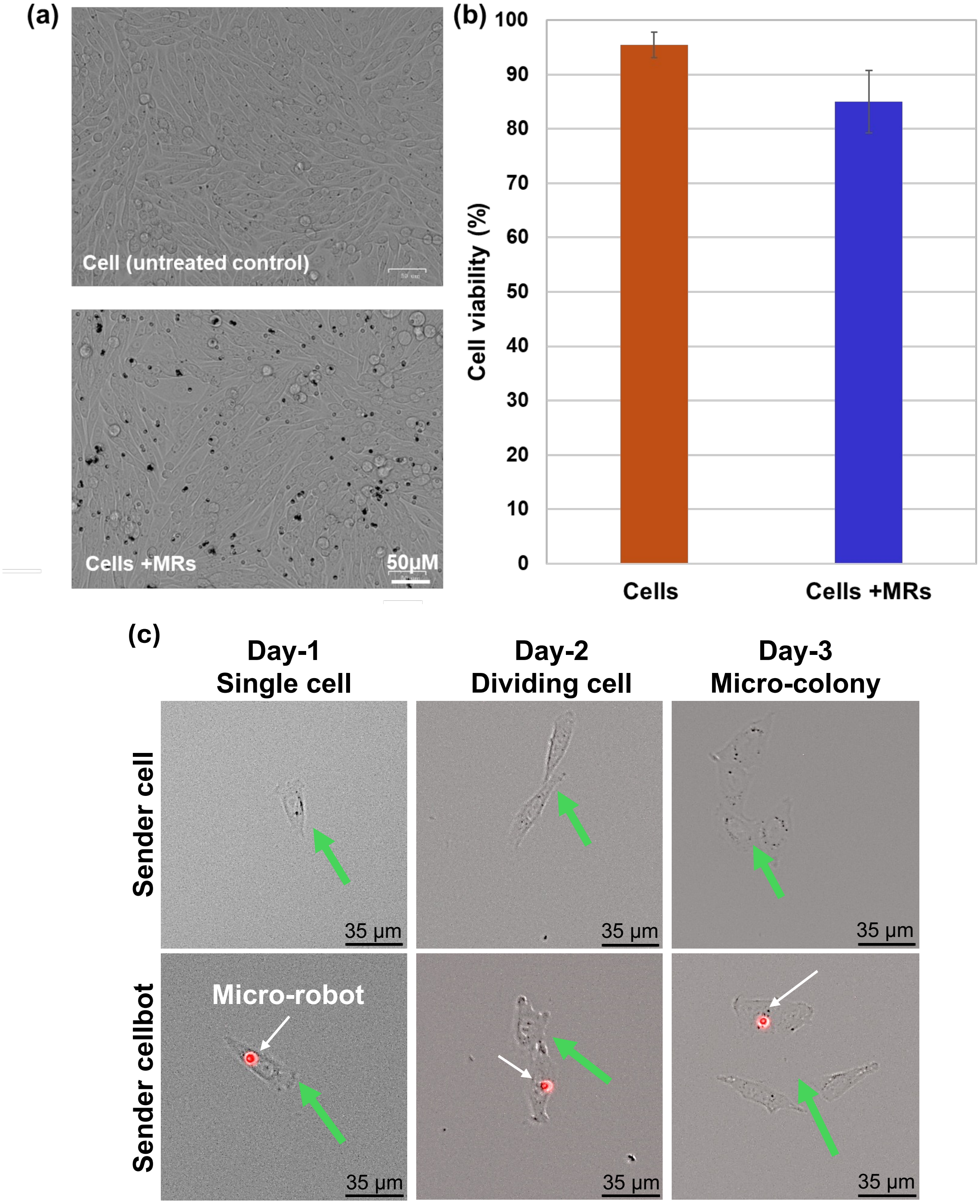


Figure SI7: **(a)** Brightfield images of control sender cell and sender cellbot cultures. **(b)** Quantitative analysis of cell viability after MRs incubation for 24hrs. **(c)** Brightfield and fluorescence images of sender cells and sender cellbots dividing over the course of 3 days.

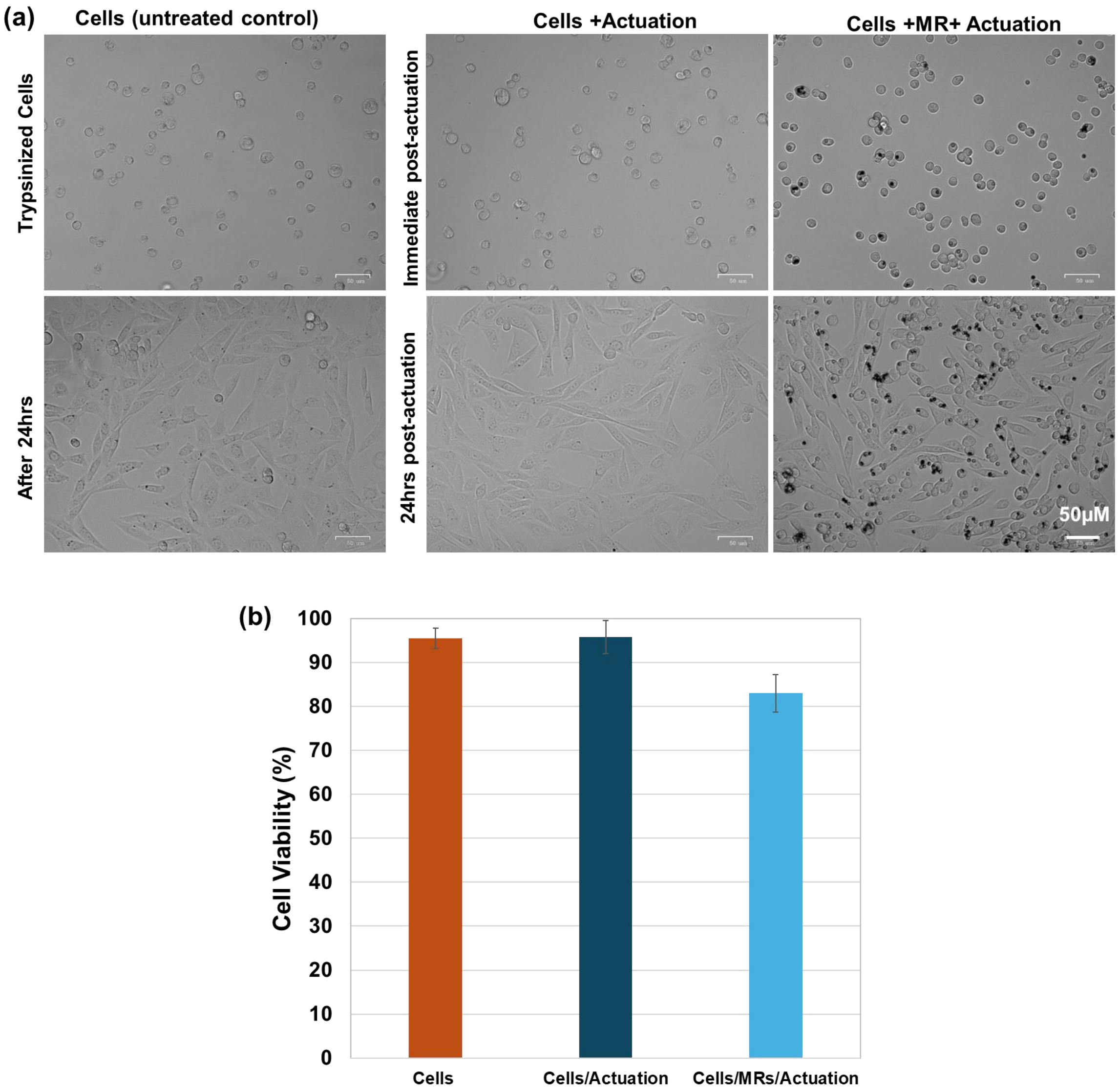


Figure SI8: **(a)** Microscopic images showing morphology and proliferation of control sender cells without MR incubation (untreated control), sender cells (cells+Actuation), and sender cellbots (cells+MRs+Actuation) immediatetly and 24 hours post magnetic actuation. **(b)** Comparison of cell viability of untreated control cells, and sender cells and sender cellbots after magnetic actuation.

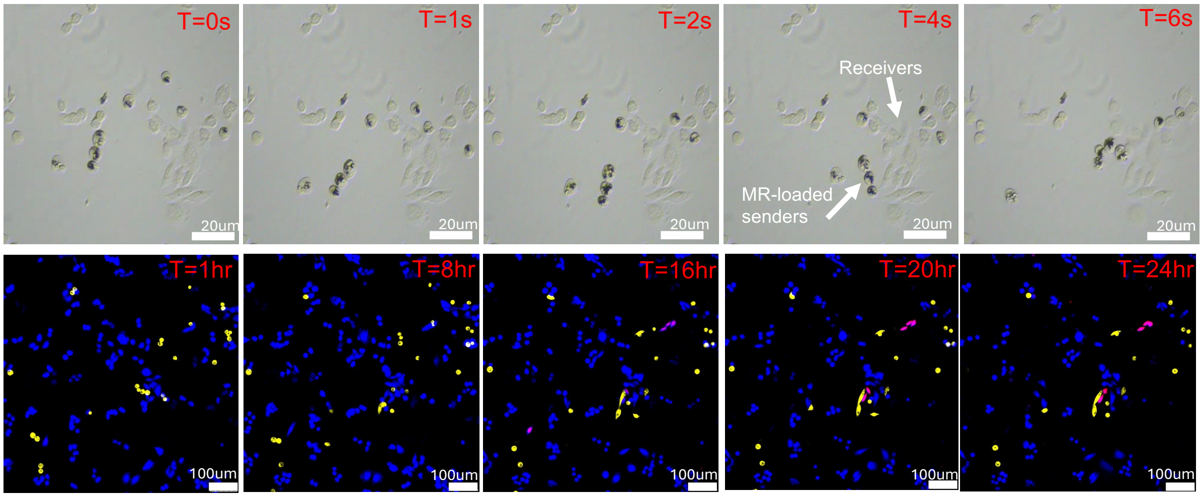


Figure SI9: MR-guided synNotch activation in CHO cells. Sender cellbots were guided to target receiver cells in a CHO receiver culture. Time-lapse images show synNotch activation when the senders cellbots maintained contact with target receivers

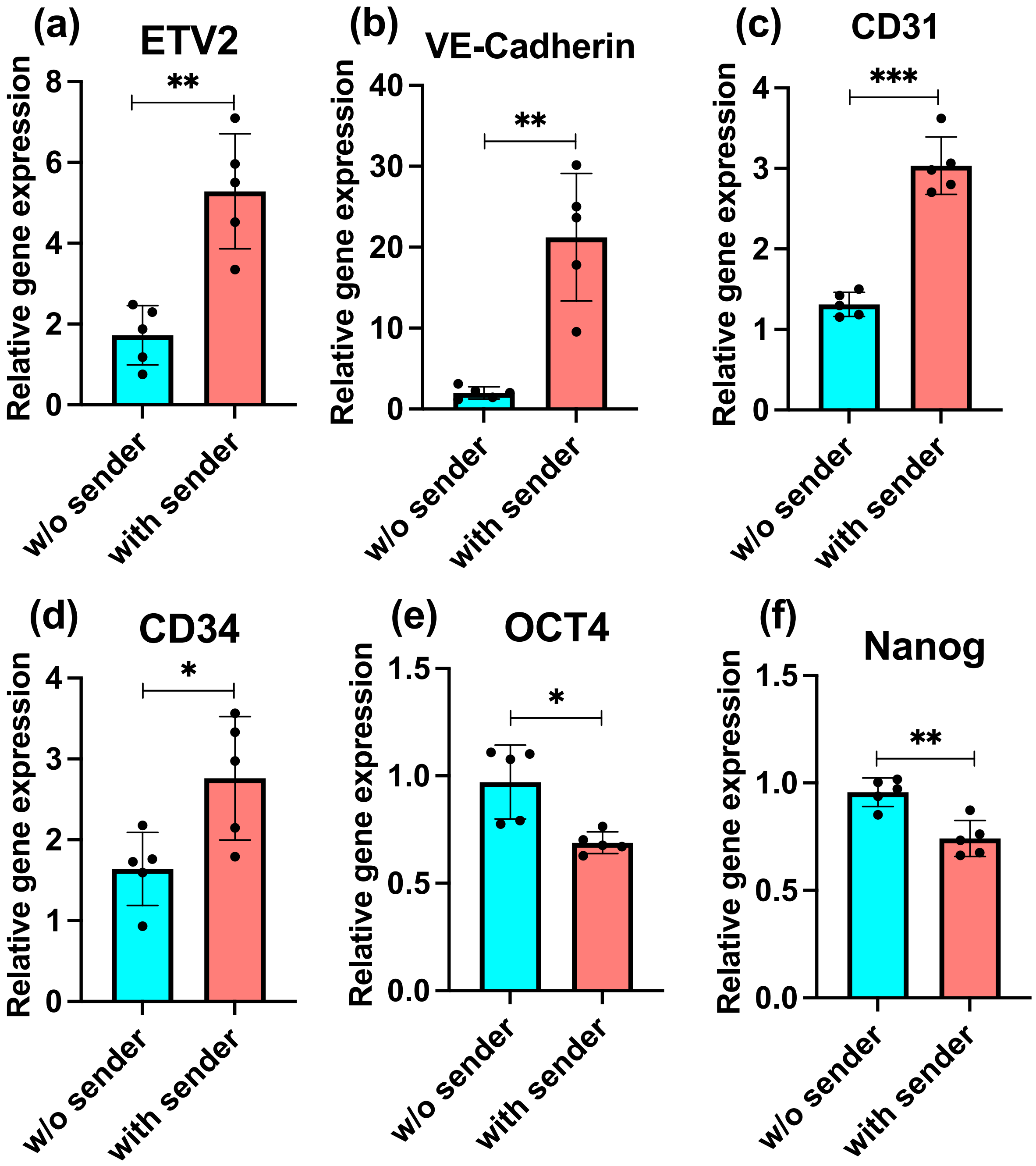


Figure SI10: RT-qPCR verifies ETV2-expression based differentiation of hiPSCs into endothelial cells. Significant increase in expression of (a) ETV2, and EC-specific markers such as, (b) VE-Cadherin, (c) CD31, and (d) CD34, were observed after 8 days of differentiation. Expression of pluripotency markers (e) OCT4 and (f) Nanog decreased in differentiated hiPSCs.

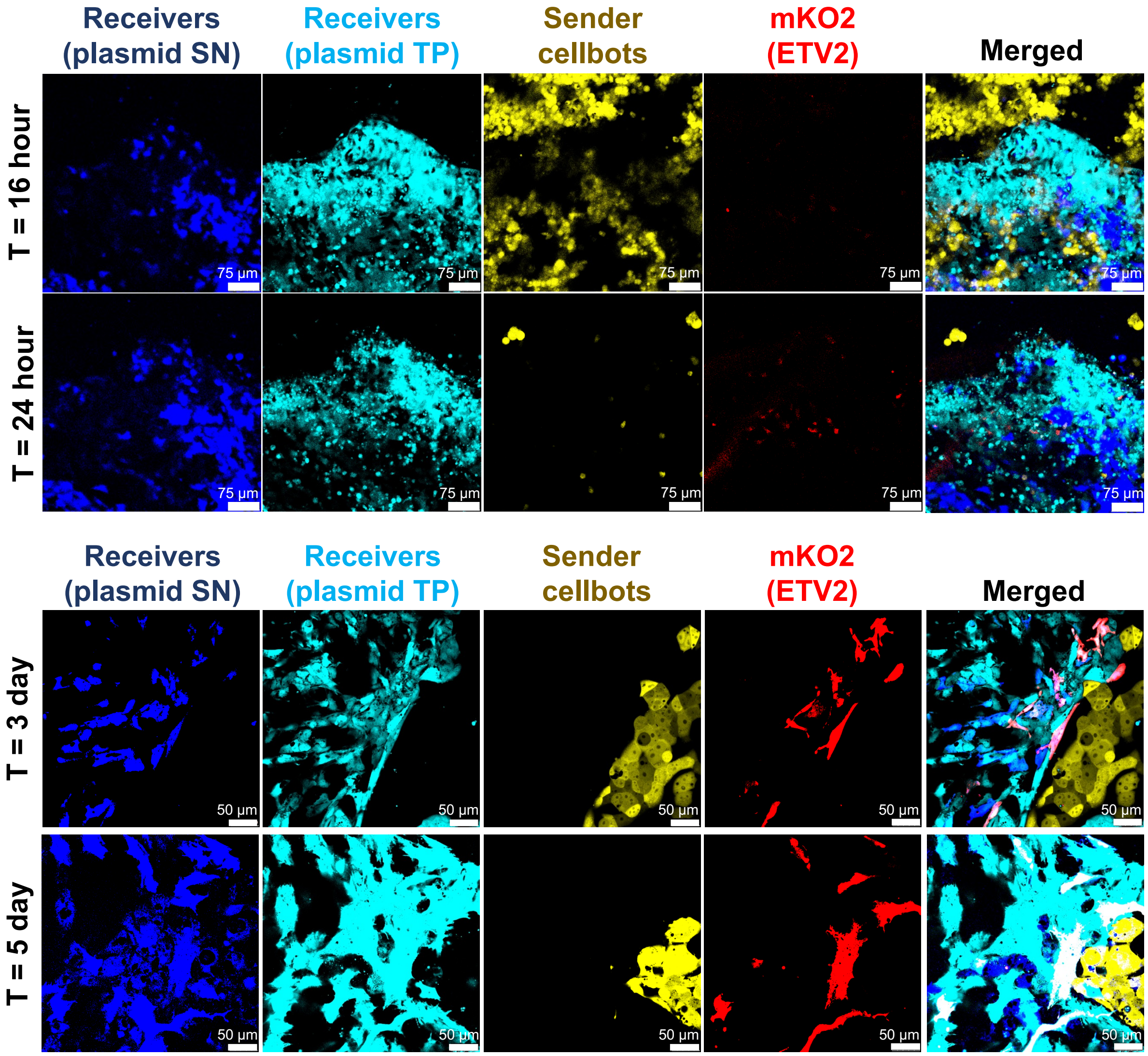


Figure SI11: (a) SynNotch activation-based mKO2 expression in hiPSC receivers in the target region after 16 and 24 hours. (b) mKO2 expression in examined targeted regions after 3 and 5 days.

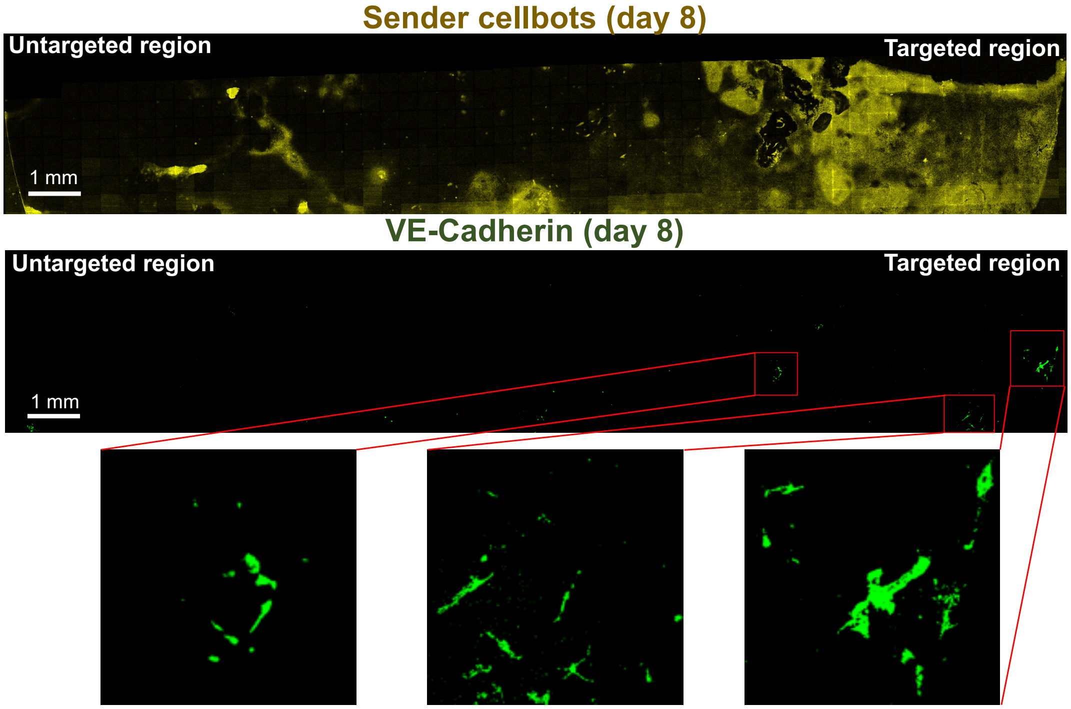


Figure SI12: Representative fluorescence images of hiPSC receivers in the target region differentiating into endothelial cells upon sender cellbots guidance to the target region. Images are acquired from two experimental repeats.

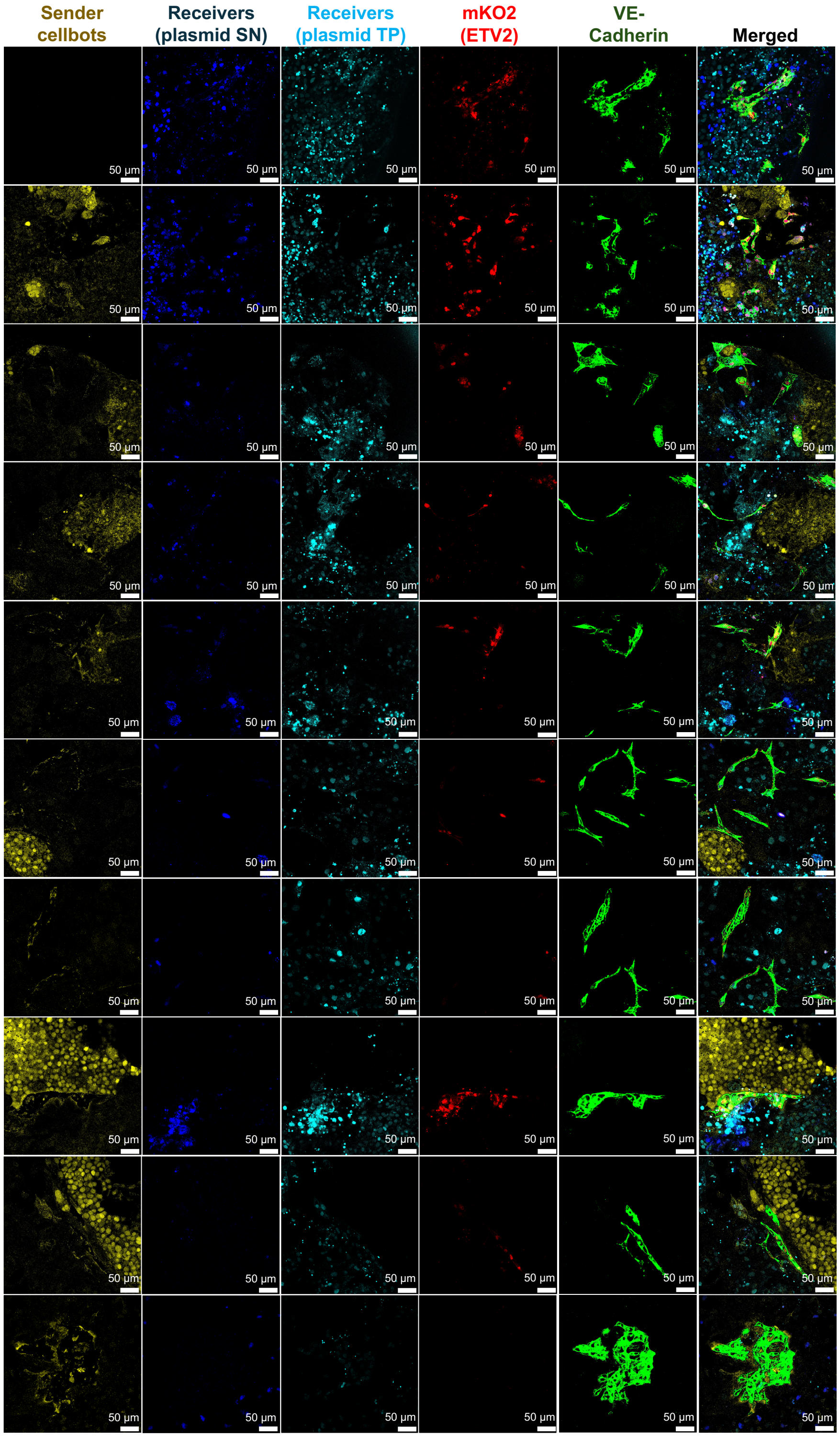

Figure SI13: Representative fluorescence images of hiPSC receivers in the target region differentiating into endothelial cells upon sender cellbots guidance to the target region. Images are acquired from two experimental repeats.

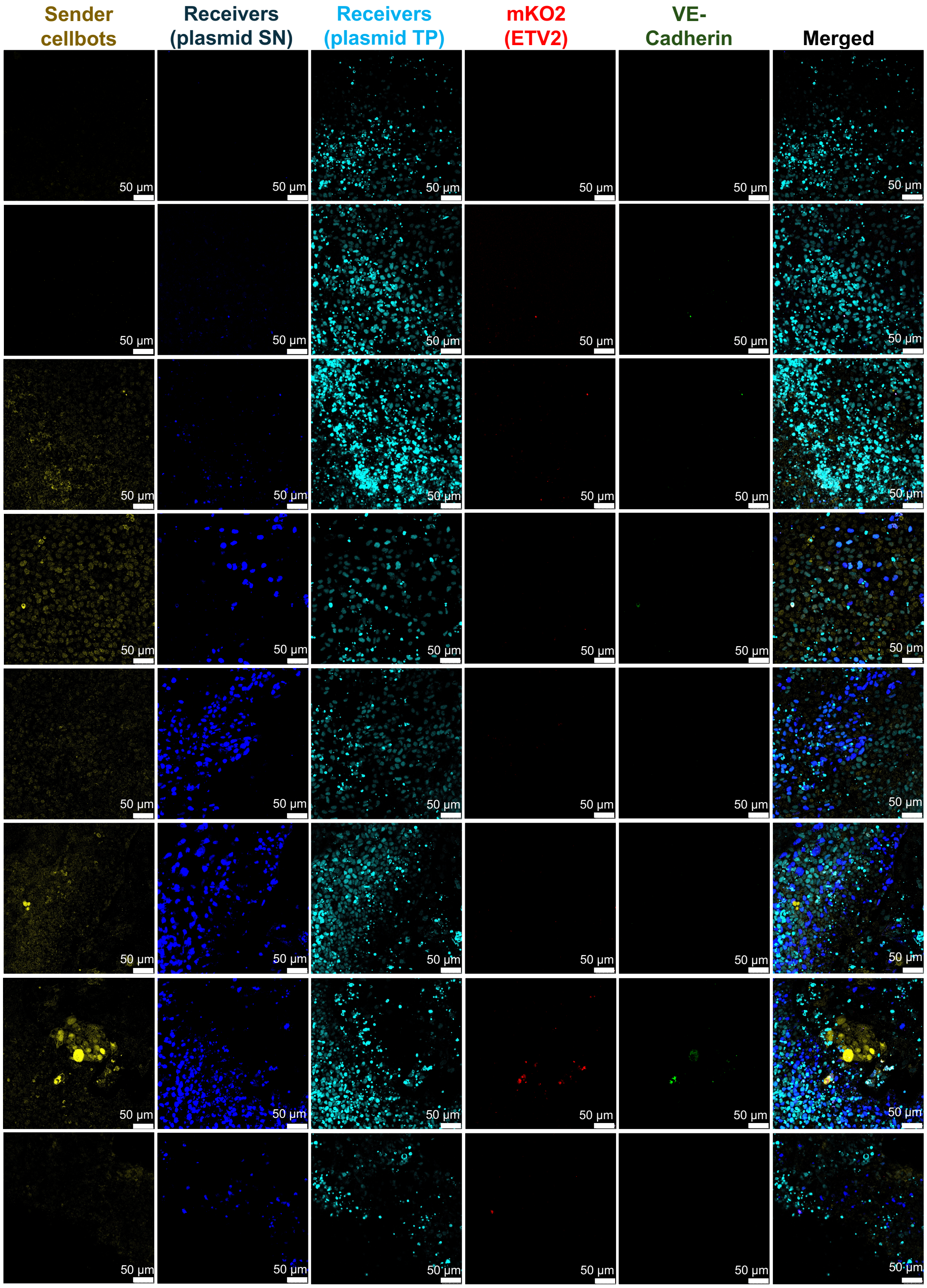


Figure SI14: Representative fluorescence images of hiPSC receivers in the untargeted region showing absence of hiPSC differentiation.

**Table SI2: Genetic circuits used to engineer synNotch sender and receiver cells**
Multiple transcriptional units (TUs) were placed in tandem orientation on the same vector backbone. The double colon (::) is used to separate the different TUs.

| Circuit ID | Circuit description |
|---|---|
| pKK499 | hEF1$\alpha$–memCD19 |
| pKK493 | hEF1$\alpha$–$\alpha$CD19-Notch core-tTA |
| pKK562 | TRE–mKate |
| pKK589 | hEF1$\alpha$–EBFP2 |
| pRM105 | TREtight–ETV2-P2A-mKO2::hEF1$\alpha$–iRFP720 |
| pRM106 | hEF1$\alpha$–$\alpha$CD19-Notch core-tTA::hEF1$\alpha$–EBFP2 |

**Table SI3: Primers used for RT-qPCR**

| Transcripts | Forward primer | Reverse primer |
|---|---|---|
| GAPDH | GGTGGTCTCCTCTGACTTCAACGT | GTGGTCGTTGAGGGCAATG |
| ETV2 | AGGGAACAAGCTGGCAGGGCTTGAA | TCCAGCATGTCTCTGCTGTCGCTGT |
| VE-Cadherin | TGTGGGCTCTCTGTTTGTTG | CGACGATGAAGCTGTATTGC |
| CD31 | GAGTATTACTGCACAGCCTTCA | AACCACTGCAATAAGTCCTTTC |
| CD34 | CTACAACACCTAGTACCCTTGGA | GGTGAACACTGTGCTGATTACA |
| OCT4 | CTTGAATCCCGAATGGAAAGGG | GTGTATATCCCAGGGTGATCCTC |
| Nanog | AGAAGGCCTCAGCACCTAC | GGCCTGATTGTTCCAGGATT |

### 3 References


[1] Lidong Yang, Yabin Zhang, Chi-Ian Vong, and Li Zhang. Automated control of multifunctional magnetic spores using fluorescence imaging for microrobotic cargo delivery. In *2018 IEEE/RSJ International Conference on Intelligent Robots and Systems (IROS)*, pages 1–6. IEEE, 2018.

[2] Chengwei Ye, Jia Liu, Xinyu Wu, Ben Wang, Li Zhang, Yuanyi Zheng, and Tiantian Xu. Hydrophobicity influence on swimming performance of magnetically driven miniature helical swimmers. *Micromachines*, 10(3):175, 2019.
